# Estrogen Receptor α/14-3-3 molecular glues as alternative treatment strategy for endocrine resistant breast cancer

**DOI:** 10.1101/2024.04.25.591105

**Authors:** Emira J. Visser, Maria Donaldson Collier, Joseph C. Siefert, Markella Konstantinidou, Susana N. Paul, Jari B. Berkhout, Johanna M. Virta, Bente A. Somsen, Peter Cossar, Galen Miley, Lara Luzietti, Leonie Young, Damir Vareslija, Lakjaya Buluwela, Simak Ali, Onno C. Meijer, Michelle R. Arkin, Christian Ottmann, Wilbert Zwart, Luc Brunsveld

## Abstract

Endocrine resistance in breast cancer treatment is a major clinical hurdle, causing an urgent need for alternative treatment modalities. The suppressive protein-protein interaction (PPI) between Estrogen Receptor alpha (ERα) and the adaptor protein 14-3-3 offers such a strategy. Here, we report the biological impact of small-molecule ‘molecular glues’ of this ERα/14-3-3 PPI by using both fusicoccin-derived semi-synthetic natural products and fully synthetic covalent drug-like molecules. We show that the ERα/14-3-3 PPI is stabilized by both the natural- and synthetic glues, resulting in a suppression of ERα transcriptional activity and a blockade of breast cancer cell proliferation, both in cell lines and in organoids derived from endocrine therapy resistant breast cancer patients. Importantly, the molecular glues effectively blocked ERα action even in case of constitutively active clinical ERα mutations, providing the foundations for developing alternative classes of ERα targeting compounds to improve treatment of patients with endocrine-therapy resistant breast cancer.

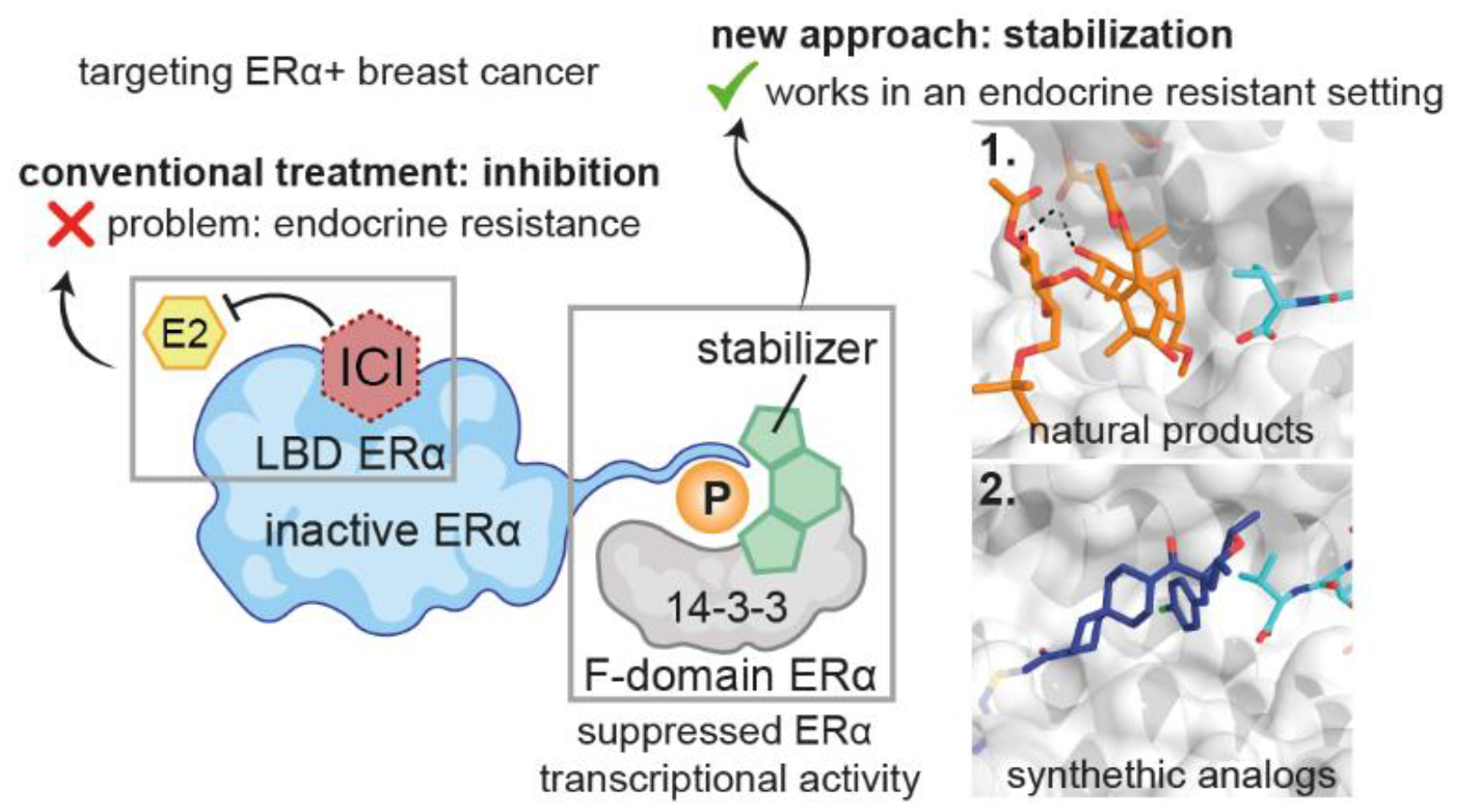

## Introduction

Breast cancer is the most common form of cancer in women and nearly 80% of these tumors express the Estrogen Receptor alpha (ERα).^1^ ERα (encoded by gene *ESR1*) is a nuclear receptor that is activated by the sex hormone estrogen and is critical to the development for this large group of breast cancers.^2^ Therefore, inhibition of ERα estrogen binding or suppressing estrogen production, both abrogating ERα-coregulator recruitment, is currently the standard of care for the treatment of hormonal breast cancer. This so-called endocrine therapy includes drugs such as Selective Estrogen Receptor Modulators (SERMs, e.g. tamoxifen), Selective Estrogen Receptor Degraders (SERDs, e.g. fulvestrant), and Aromatase Inhibitors (AIs, e.g. letrozole).^3^ While these treatments substantially reduce cancer recurrence and mortality, 20% of patients do recur with ERα positive, endocrine-resistant Metastatic Breast Cancer (MBC)^4^, thus presenting a major clinical hurdle. The mechanisms of endocrine resistance are extensively studied^5^, including ligand-independent ERα reactivation through point-mutations in the ligand-binding domain (LBD) of ESR1,^6^ most commonly at Tyr537 and Asp538. These mutations allow for ligand-independent ERα reactivation, leading to resistance to AIs and reduced sensitivity to tamoxifen and fulvestrant.^7–10^ As such, there is a pressing need for the development of alternative, molecularly orthogonal therapeutics, preferably targeting ERα outside its LBD, for treating endocrine resistant MBC.^11^

An appealing negative regulator of ERα is the phosphoserine/threonine-binding adaptor protein 14-3-3. The seven isoforms of 14-3-3 have highly overlapping functions in many pathophysiological pathways, including cancer.^12^ Phosphorylation of ERα at its penultimate residue Thr594, outside the LBD, enhances binding to all isoforms of 14-3-3, resulting in reduced estradiol-dependent transcriptional activity.^13^ 14-3-3 serves as an endogenous ERα interactor, as reported in Rapid Immunoprecipitation Mass Spec of Endogenous proteins (RIME) experiments performed by different groups over the last decade^14–17^, suggesting a hormone-independent strategy to target ERα positive breast cancer by stabilizing the ERα/14-3-3 complex. This unique mechanism can be enhanced by the natural product Fusicoccin-A (FC-A), which acts as a “molecular glue” by binding at the ERα/14-3-3 interface.^13^ The development of such PPI stabilizers is an emerging field in drug discovery^18,19^, exemplified by the development of molecular glue degraders and PROteolysis TArgeting Chimeras (PROTACs).^20–22^

While the natural product FC-A is a useful tool to validate the ERα/14-3-3 molecular glue concept, its clinical utility is limited by its complex structure which limits diversity via (semi)-synthesis methodologies^23^, by the difficulty in isolating it from natural sources^24^, and by the high micromolecular concentrations needed to achieve a partially effective dose.^13^ Therefore, we developed alternative small-molecule stabilizers of the ERα/14-3-3 PPI.^25^ These synthetic stabilizers target a native cysteine residue on 14-3-3σ and bind in the same composite ERα/14-3-3 binding pocket, also occupied by FC-A.^25^

Here, we investigate the biological mechanisms of a range of fusicoccin derivatives and novel synthetic stabilizers to fully grasp the underlying biological consequences of ERα/14-3-3 PPI stabilization and ultimately to develop an alternative therapeutic strategy to target endocrine resistant breast cancer. We first investigate the stabilization potential of the natural product FC-A and six of its semi-synthetic derivatives by performing binding studies, crystallizing the ternary complexes (ERα-peptide/14-3-3/stabilizer), and evaluating cell proliferation. FC-A suppressed ERα transcriptional activity in MCF-7 breast cancer cells. The improved stabilizer FC-NAc and synthetic covalent ERα/14-3-3 stabilizers also suppressed ERα activity in MCF-7 cells, however at significantly lower effective concentrations than FC-A. Mechanistically, decreased ERα transcriptional activity coincided with increased ERα/14-3-3 interactions, as observed by Proximity Ligation Assays (PLA). Importantly, ERα/14-3-3 stabilization was still efficacious for constitutively active mutations ERα (Y537S, D538G) and in endocrine-resistant human organoid models. Decreased cell proliferation capacity following ERα/14-3-3 PPI stabilization also acted synergistically with fulvestrant (ICI 182780; “ICI”) treatment, further suppressing cellular and organoid proliferation. Our data supports the inhibition of ERα in MBC using this novel molecular-glue strategy in combination with existing endocrine therapeutics.

## Results

### FC-NAc is the most effective FC-A-derived stabilizer of the ERα/14-3-3 complex

ERα/14-3-3 PPI glues were identified by testing seven semi-synthetic derivatives of FC-A for their stabilization potential (Fig. S1a). This focused library included the 3’-deacetylated analogue of FC-A (deAc-FC-A), fusicoccin-J (FC-J)^26^, FC-J acetonide (FC-J-Ace)^27^, FC-31 that contains an introduced piperidine ring, FC-THF^28^ with its bulky tetrahydrofuran substitution on ring C, FC-NAc^29^ with an acetamide group exchanged for the 9-acetoxy group of FC-A, and lastly the aglycon derivative of FC-NAc, FC-NAg. Using Fluorescence Anisotropy (FA), we measured the half-maximal effective concentration (EC_50_), of the compounds to stabilize the interaction between 14-3-3σ and a representative C-terminal ERα peptide labeled with fluorescein (FAM). The EC_50_ values were in the nanomolar range for all fusicoccanes except for FC-THF and FC-NAg, which exhibited an EC_50_ value in the micromolar range (Fig. S1b). The FC-NAc derivative showed the greatest stabilization efficiency (EC_50_ = 15 nM) compared to FC-A (EC_50_ = 421 nM) and FC-NAg showed the weakest (Fig. 1a, 1b).

**Fig. 1.**
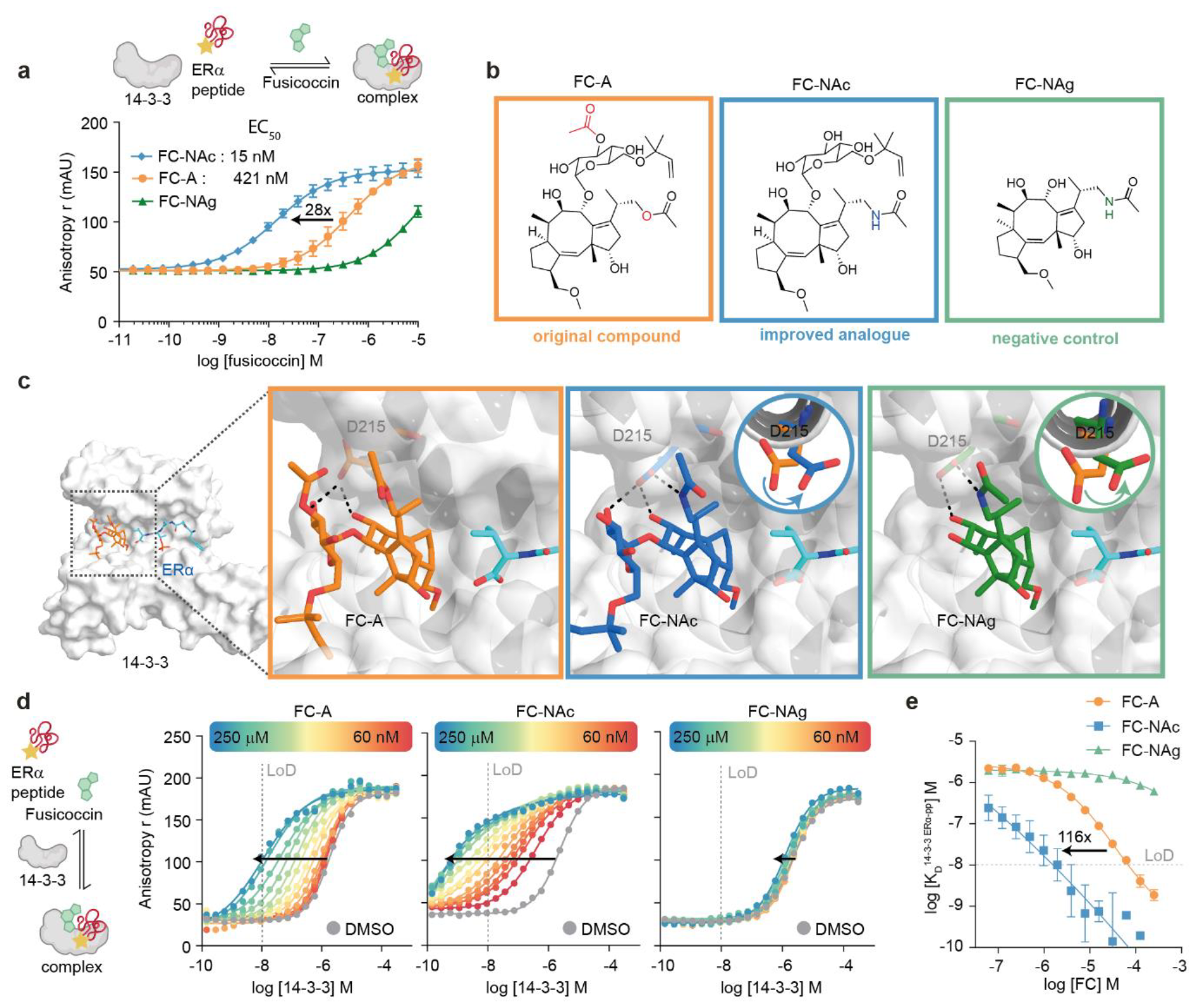
FC-NAc is an improved stabilizer of the ERα/14-3-3 PPI interaction. **a**, Fluorescence Anisotropy (FA) assay with titration of FC-A (orange), FC-NAc (blue) and FC-NAg (green) to FAM-labeled ERα peptide (10 nM) and 14-3-3σ (1 μM) (mean ± SD, n = 3). **b**, Molecular structures of FC-A, FC-NAc and FC-NAg. **c**, Crystal structures of 14-3-3 (white surface), phospho-ERα peptide (cyan sticks), and FC-A (left, orange sticks), FC-NAc (middle, blue sticks) and FC-NAg (right, green sticks). **d**, FA 2D titrations with titration of 14-3-3σ to FAM-labeled ERα peptide (10 nM), against varying concentrations (0– 250 μM) of FC-A (left), FC-NAc (middle) and FC-NAg (right). **e**, Plot of the apparent K_D_ values derived from the 14-3-3 titrations with varying concentrations of FC, with FC-A (orange circles), FC-NAc (blue diamonds) and FC-NAg (green triangles) (mean ± SD, n = 3).

To structurally understand the difference in EC_50_ values, the FC derivatives were further evaluated by X-ray crystallography. All derivatives showed a clear electron density and bound at the ERα/14-3-3 PPI interface (Fig. S2). In general, the best stabilizers induced 14-3-3 helix-9 to move towards ERα, effectively ‘clamping’ the complex into a closed conformation and caused the C-terminal valine of ERα to be directed towards the FCs/fusicoccanes (Fig. S3). In addition to the clamping effect observed for the best stabilizer, FC-NAc, its introduced N-acetamide also resulted in a turn of the aspartic acid residue (Asp215) of 14-3-3, thereby facilitating an additional hydrogen bond (Fig. 1c). The poor stabilization activity of the aglycone variant FC-NAg is in line with previous research on aglycone fusicoccanes^28^, and can be explained by the loss of a hydrogen bond to 14-3-3 and exposure of the hydrophobic FC core to the solvent, resulting in energetically unfavorable interactions combined with a reduced molecular surface interaction with 14-3-3.

The stabilization efficiencies of FC-NAc, FC-A, and FC-NAg were further evaluated by two-dimensional (2D) FA titrations, in which both the compound and the 14-3-3 protein concentrations were varied. Such assays evaluate the fold-decrease in the dissociation constant for the ERα/14-3-3 complex (apparent K_D_) and the intrinsic binding of the FC analog to 14-3-3 (K_D2_).^27^ Hence, 14-3-3σ was titrated to FAM labeled ERα-peptide (10 nM), in the presence of a range of concentrations of FC-A, FC-NAc or FC-NAg (0–250 μM) (Fig. 1d). Plotting the change in apparent K_D_ of the ERα/14-3-3 interaction in the presence of different stabilizer concentrations (Fig. 1e) demonstrated a 116-fold stronger decrease in EC_50_ for FC-NAc compared to FC-A. To date, FC-NAc is the most potent ERα/14-3-3 PPI natural-product-derived stabilizer reported, making it an ideal candidate for chemical biology studies into the underlying biomolecular effects of the ERα/14-3-3 interaction. For further studies, FC-NAc cellular activity was compared to the original natural product (FC-A), while the weak stabilizer FC-NAg served as a negative control.

### FC-NAc suppresses ERα transcriptional activity

The three fusicoccanes (the parental FC-A, the enhanced FC-NAc and negative control FC-NAg) were tested for their effect on cell viability of MCF-7 (ERα positive) and MDA-MB-231 (ERα negative) breast cancer cells by performing cell titer blue (CTB) assays (Fig. S4a). FC-A treatment showed an IC_50_ value of 50 μM in MCF-7 cells, confirming our previous observations^13^, while the improved stabilizer FC-NAc had an enhanced IC_50_ value of 20 μM. Neither fusicoccane impaired the viability of the ERα-negative cell line MDA-MB-231. Our negative control, FC-NAg, did not show any effect on cell viability in either cell line. Based on these results, we decided to use a concentration of 30 μM of the fusicoccanes for the remaining assays. We confirmed cell proliferation results using live-cell imaging analysis (Fig. 2a). Here, 30 μM of FC-NAc was more effective in inhibiting estradiol (E2)-induced cell proliferation of MCF-7 cells compared to FC-A. This trend was also visible in two other ERα-positive breast cancer cell lines, T47D and ZR-75-01 (Fig. S4b), qualitatively confirming our findings. Again, no effect was visible for the ERα-negative breast cancer cell line MDA-MB-231 (Fig. S4b).

**Fig. 2.**
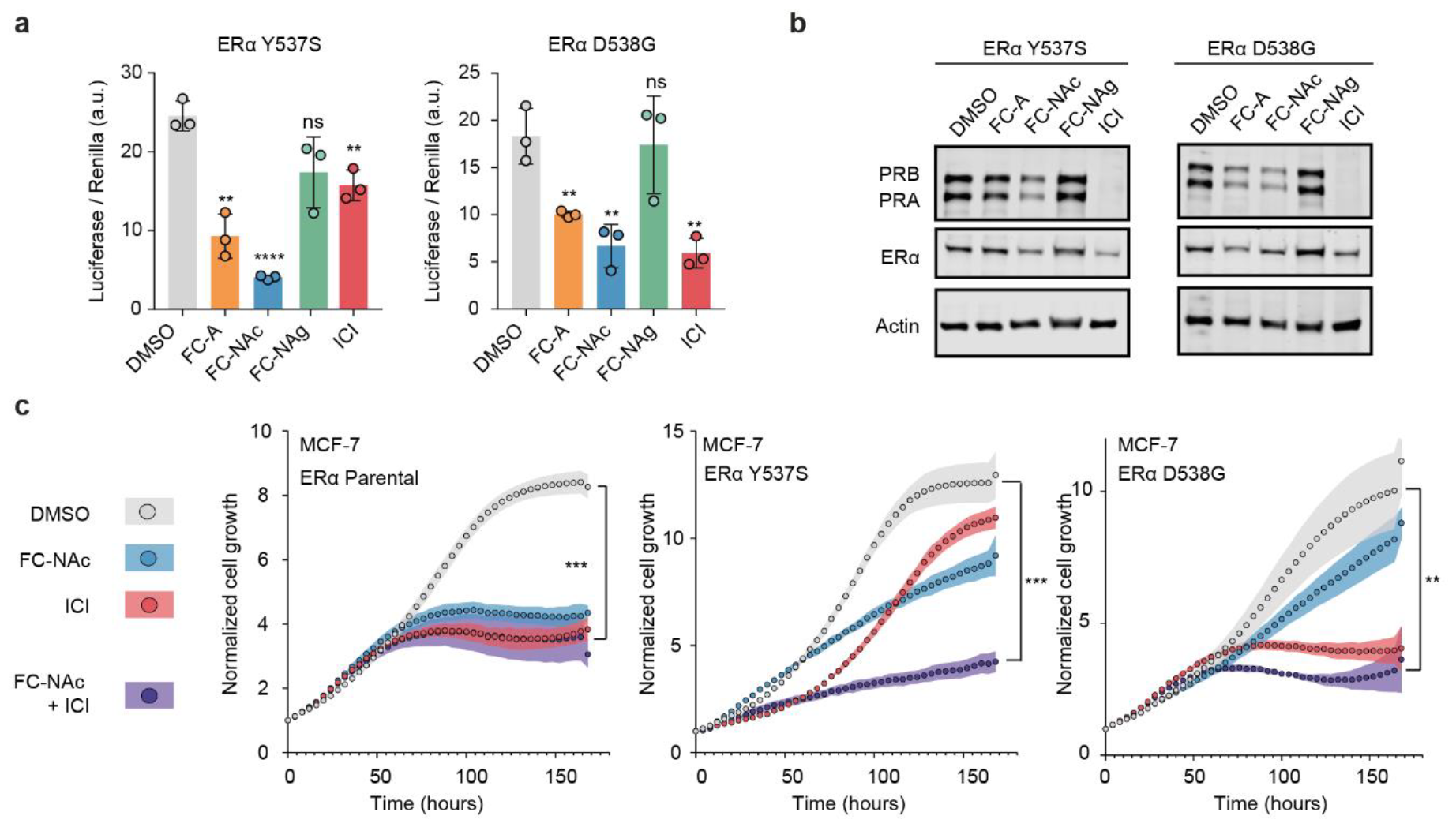
Targeting of ERα Y537S and D538G somatic mutations by the fusicoccanes. **a**, Luciferase reporter assay of ERα Y537S (left) and D538G (right) cells, transfected with ERE-luc and renilla after pretreatment with DMSO (grey bars), FC-A (orange, 30 μM), FC-NAc (blue, 30 μM), FC-NAg (green, 30 μM) or ICI (red, 100 nM). (mean ± SD, n=3 biologically independent, with each 2 technical replicates). **b**, Western blot of down-stream protein PR of the ERα pathway for MCF-7 Y537S cells (left) and MCF-7 D538G cells (right), treated with FC-A, FC-NAc, FC-NAg (all 30 μM) or ICI (100 nM) (n = 3). **c**, MCF-7 ERα WT (left), Y537S (middle) and D538G (right) cell proliferation in the presence of DMSO (grey circles), FC-NAc (blue, 30 μM), ICI (red, 100 nM) or FC-NAc (30 μM) and ICI (100 nM) combined (purple) (mean ± SD, n=3 biologically independent, with each 2 technical replicates). (unpaired t-test: **p≤0.005 ***p≤0.001, ****p≤0.0001).

Luciferase reporter assays were then performed to evaluate inhibition of ERα transcriptional activity. An ERα-driven estrogen response element-luciferase reporter (ERE-luc) plasmid was transfected into hormone-deprived MCF-7 cells, together with a plasmid containing constitutively expressed Renilla luciferase to correct for transfection efficiency. The cells were pretreated overnight with the fusicoccanes (30 μM), or fulvestrant (ICI; 100 nM) as positive control. The luciferase signal was measured after 24 hours of stimulation with increasing concentrations of E2 (Fig. 2b). At 10 nM E2, 30 μM FC-A and 30 μM FC-NAc reduced the ERα transcriptional activity by ∼47% and ∼65%, respectively, similar to 100 nM ICI. As expected, the weak stabilizer FC-NAg did not impact ERα transcriptional activity. The improved stabilization potency of FC-NAc thus results in an enhanced inhibitory effect on breast cancer cellular proliferation and inhibition of the transcriptional activity of ERα in MCF-7 cells. These results were additionally validated at the protein level using the classical and clinically-informative endogenous ERα-target gene, the progesterone receptor (PR)^30^ (Fig. 2c, Fig. S5).

ERα Thr594 phosphorylation is crucial for 14-3-3 binding, and stabilization of this PPI results in increased Thr594-ERα phosphorylation levels.^13^ Both FC-A and FC-NAc increased Thr594 phosphorylation of ERα, in contrast to ICI (Fig. 2d, Fig. S6). To confirm our prior observations^13^, we treated MCF-7 cells with the proteasome inhibitor MG-132 (5 μM) a day before cell harvesting; inhibition of the proteasome slightly increased Thr594 phosphorylation following FC-A and FC-NAc, commensurate with the increased ERα levels in the cell.

To measure impact on ERα transcriptional activity on a global level, RNA-sequencing was performed on MCF-7 cells treated with the fusicoccanes (FC-A, FC-Nac, FC-Nag, all 30 μM) for three days followed by E2 (10 nM) stimulation for six hours. Principal Component Analysis (PCA) of the RNA-seq results indicated that FC-A and FC-NAc treatment differed distinctly from the DMSO and FC-NAg treatment in their gene expression profiles (Fig. S7). Differentially Expressed Genes (DEGs) were identified for FC-NAc (509 genes), FC-A (418 genes) and negative control FC-NAg (163 genes) (Fig. 2e, Extended Data Table 1), with 203 DEGs specifically shared between FC-NAc and FC-A (Fig. 2f). A Gene Set Over-representation Analysis (GSOA) showed a suppression of the early and late estrogen response for both the FC-A and FC-NAc treatment (Fig. 2g).

**Fig. 1.**
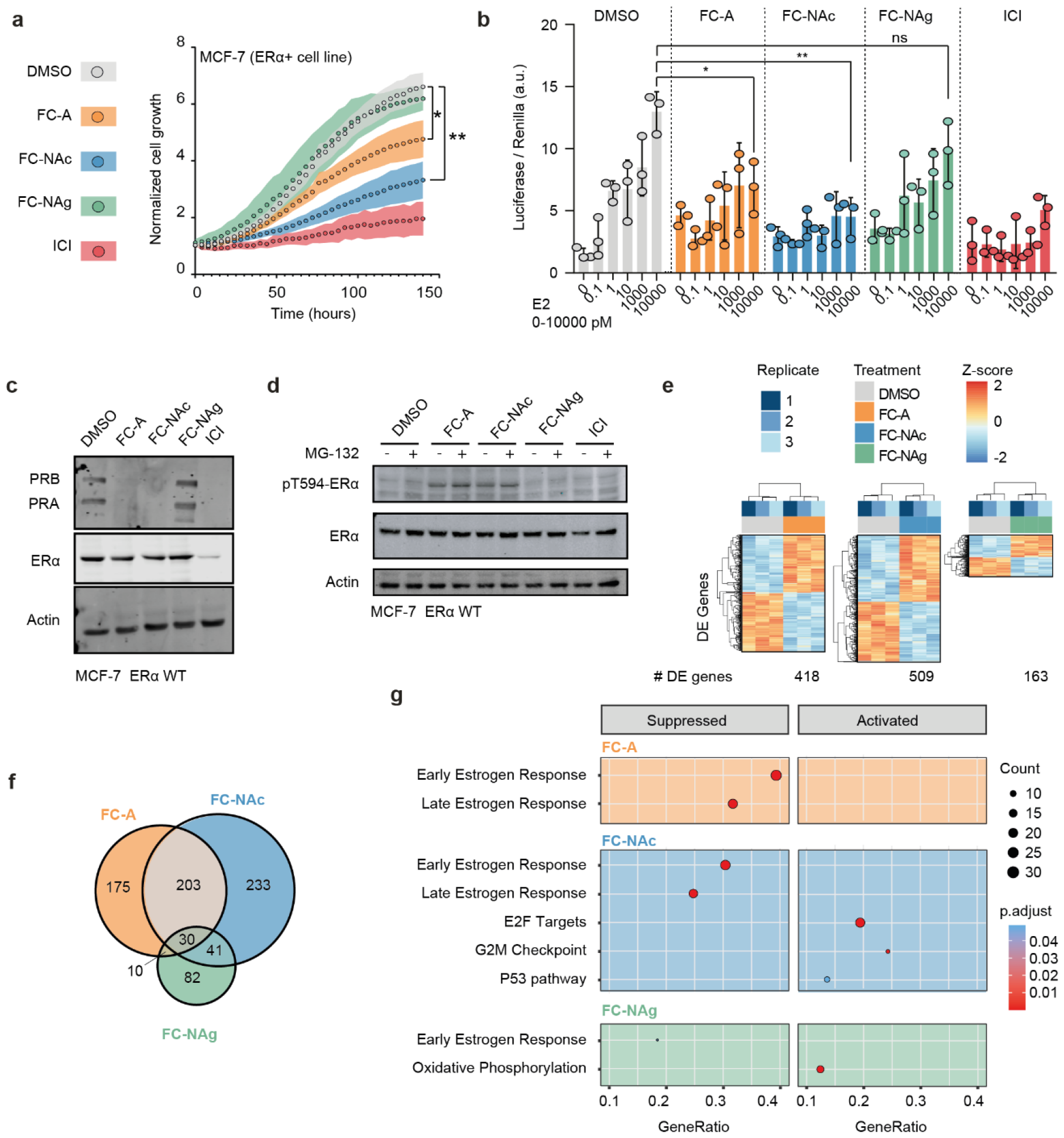
FC-NAc suppresses ERα transcriptional activity. **a**, E2-induced MCF-7 cell proliferation in the presence of DMSO (grey), FC-A (orange, 30 μM), FC-NAc (blue, 30 μM), FC-NAg (green, 30 μM) or ICI (red, 100 nM). (mean ± SD, n=3 biologically independent, each 3 technical replicates). **b**, Luciferase reporter assay of MCF-7 cells transfected with ERE-luc and renilla for various E2 concentrations, pretreated with DMSO (grey), FC-A (orange, 30 μM), FC-NAc (blue, 30 μM), FC-NAg (green, 30 μM) or ICI (red, 100 nM) (mean ± SD, n=3 biologically independent, with each 2 technical replicates). **c**, Western blot of down-stream protein PR of the ERα pathway for MCF-7 cells treated with FC-A, FC-NAc, FC-NAg (all 30 μM) or ICI (100 nM). **d**, Western blot of pT594-ERα phosphorylation levels in MCF-7 cells treated with DMSO, FC-A, FC-NAc, FC-NAg (all 30 μM) or ICI (100 nM), with and without treatment of MG-132 (10 μM). **e**, Heat-map depicting the changes in gene expression after FC-A (left), FC-NAc (middle) or FC-NAg (right) treatment. **f**, Number of shared and unique differentially expressed genes with FC-A (orange), FC-NAc (blue) or FC-NAg (green) treatment. **g**, Gene set over-representation analysis (GSOA) of differentially expressed genes upon treatment of FC-A (top, orange), FC-NAc (middle, blue), or FC-NAg (bottom, green). (Unpaired t-test, *p≤0.05, **p≤0.005).

### Fusicoccanes retain activity against ERα with somatic mutations Y537S and D538G

The most common mutations in the ERα gene, which have been linked to acquired resistance in endocrine therapies, are Y357S and D538G.^7^ Cell studies demonstrated that these activating mutations required higher concentrations of antagonist to inhibit ERα signaling compared to the wild-type (WT) receptor.^31^ As these mutations are located near the LBD of ERα, we hypothesized that they would not hamper ERα/14-3-3 stabilization, which occurs at the C-terminal ERα F-domain; 14-3-3 stabilization might therefore offer an orthogonal strategy to inhibit these constitutively active ERα variants (Fig. S8). Two MCF-7 ESR1 Exon 8 knock-in cell-line mutations Y537S and D538G^32^ were used to study this hypothesis. Luciferase reporter assays illustrated suppression of mutant ERα activity by FC-A and FC-NAc to a similar level as the wild-type cell line (Fig. 3a), which could be confirmed at the protein level for the known ERα target, PR (Fig. 3b, Fig. S9-S13). The ERα degrader fulvestrant (ICI) also blocks ERα-mediated transcription for these mutations, although with a reduced efficiency.^33^ Interestingly, while transcriptional blockade was similar to the parental cell lines, FC-NAc and ICI inhibited cell-growth somewhat less effectively for the mutants compared to the parent (Fig. 3c); moreover, FC-NAc-mediated growth inhibition could be further enhanced by addition of ICI. Notably, the ERα D538G cell line was more sensitive to ICI treatment compared to the ERα Y537S cell line, consistent with several published studies^31,34^, in contrast to the greater sensitivity of Y537S cells to FC-NAc treatment.

**Fig. 3.**
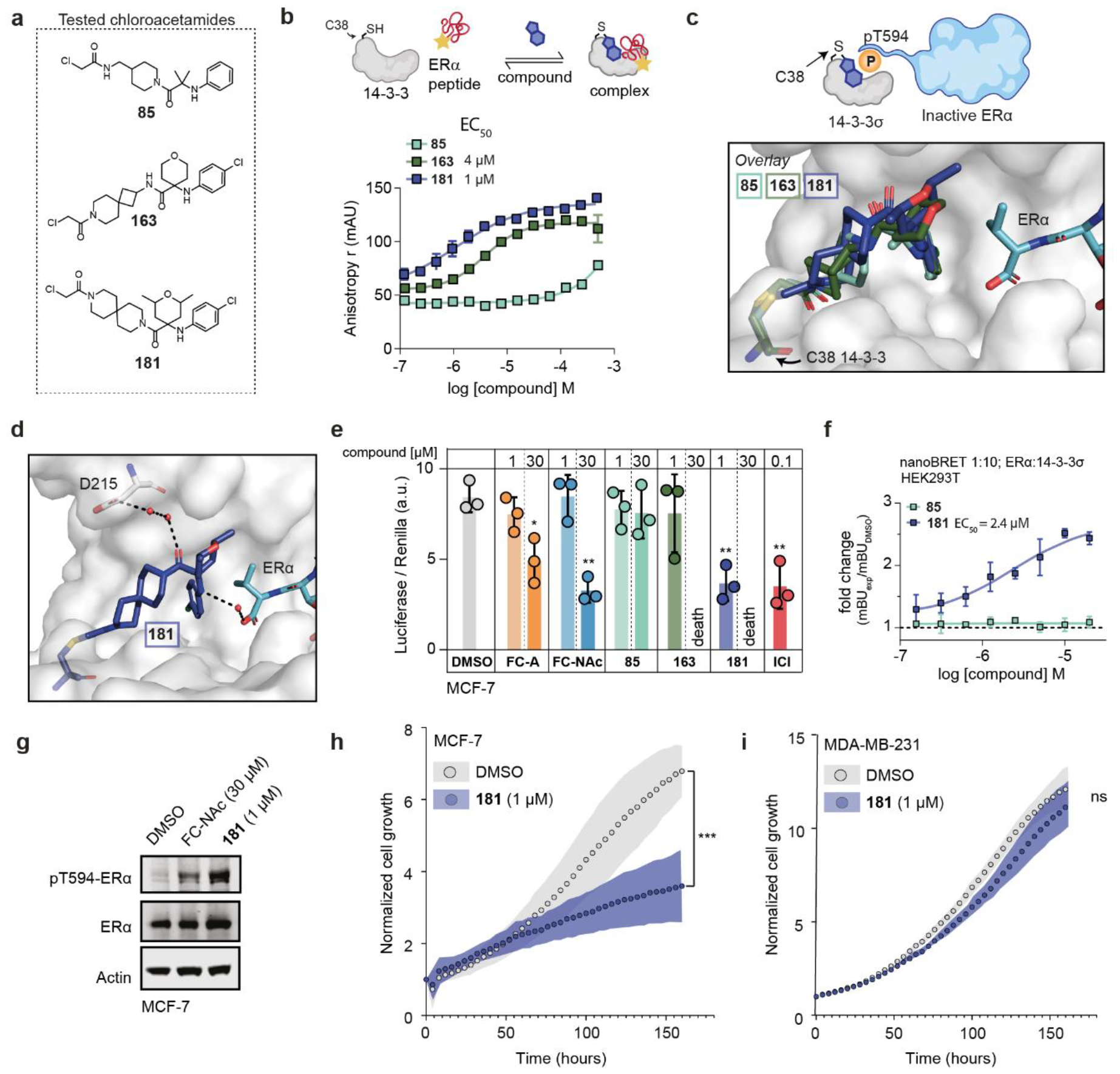
Synthetic covalent stabilizers suppress ERα activity. **a**, structure of three chloro-acetamide stabilizers tested. **b**, Fluorescence Anisotropy (FA) assay with titration of **85** (light green), **163** (dark green) and **181** (purple) to FAM-labeled ERα peptide (10 nM) and 14-3-3σ (1 μM) (mean ± SD, n = 3). **c**, crystallographic overlay of **85** (light green; PDB = 8AI0), **163** (dark green; PDB = 8AQC) and **181** (purple; PDB = 8APS) in complex with 14-3-3σ (white surface) and ERα (cyan sticks). **d**, crystal structure of **181** (purple) in complex with 14-3-3σ (white surface) and ERα (cyan sticks) (PDB = 8APS). Water molecules are represented as red spheres with its interactions as black dashes. **e**, Normalized transactivation activity (ERE-luciferase assay) of MCF-7 cells transfected with ERE-luc and Renilla in the presence of 1 μM or 30 μM of the fusicoccanes (FC-A, FC-NAc), the chloro-acetamides or 100 nM ICI as positive control (mean ± SD, n=3). **f**, NanoBRET signal observed for ERα/14-3-3σ hybridization in HEK293T cells with increasing concentration of **85** (light green) or **181** (purple) (mean ± SD, n = 2). **g**, Western blot of Thr594-ERα phosphorylation levels in MCF-7 cells upon treatment of FC-NAc (30 μM) or **181** (1 μM). **h-i**, Cellular proliferation assay with DMSO or 1 μM of **181** in MCF-7 cells (**h**) or MDA-MB-231 cells (**i**) (mean ± SD, n=3). (unpaired t-test: *p≤0.05, **p≤0.005 ***p≤0.001).

### Synthetic stabilizers suppress ERα transcriptional activity at lower effective concentrations

The promising cellular responses following ERα/14-3-3 PPI stabilization by the fusicoccanes directed our efforts to the development of fully synthetic stabilizers of the ERα/14-3-3 PPI^25^, since the complex structure of fusicoccanes hampers further medicinal chemistry optimization. Chloroacetamide-containing compounds were developed to covalently bind to Cys38 of the 14-3-3σ isoform to specifically stabilize the ERα/14-3-3σ PPI (Fig. 4a). We evaluated three novel chloro-acetamide derivatives with a range of EC_50_ values, as confirmed by Fluorescent Anisotropy (FA) studies (Fig. 4b). X-ray crystallography provided a rationale for the activity of these three analogs (Fig. 4c). Briefly, the chloroaniline moiety in compounds **163** and **181** bound into a composite pocket formed by 14-3-3 and the C-terminal V595 of ERα; similar to the binding of FC-NAc, **181** engages with the composite ERα/14-3-3 pocket via several water molecules and made a hydrogen bond with Asp215 on 14-3-3 and with the carboxylic acid of the ERα C-terminus (Fig 4d). Compound **85** lacked the phenyl-Cl group resulting in low activity in FA assays, while compounds **163** and **181** were amongst the best stabilizers, with an EC_50_ value of 4 μM and 1 μM, respectively (Fig. 4b).

Interestingly, 1 μM of **181** reduced ERα transcriptional activity to the same degree as 30 μM of FC-NAc and 100 nM of ICI, as demonstrated by ERα luciferase reporter assays (Fig. 4e). At 30 μM, **163** and **181** appeared to be cytotoxic. Biochemically inactive compound **85** also showed no effect in the luciferase reporter assay. To study the intracellular interaction between ERα and 14-3-3σ, a nanoBRET assay was designed. Briefly, Nluc-ERα:14-3-3σ-HaloTag plasmids (1:10 ratio) were transfected in HEK-293T cells, followed by hormone deprivation for 48 hours. Cells were then trypsinized, re-seeded, and treated with 100nM HaloTag ligand. After cells adhered (∼1 hour), cells were treated with chloro-acetamides **85** or **181** for 24 hours in 1:2 dilution series starting at 20 μM and the BRET signal was read and normalized against DMSO treatment. Interestingly, **181** showed an increase in ERα/14-3-3 hybridization with an EC_50_ value of 2.4 μM while for **85** no hybridization was observed (Fig. 4f), which is in accordance with the luciferase reporter assay. No increase in ERα/14-3-3 hybridization was observed when transfecting the 14-3-3σ C38N construct (Fig. S14), confirming the necessity for covalent binding to C38 by the stabilizers.

The effect of ERα/14-3-3 stabilization by the most potent synthetic stabilizer **181** was further analyzed by staining for Thr594 ERα phosphorylation levels using western blot. Molecular glue **181** increased phosphorylation levels more than 30 μM of FC-NAc (Fig. 4g, Fig. S15). This data indicated that **181** effectively enhanced binding of 14-3-3 to its ERα phospho-site, thereby protecting it from phosphatases. Further, 1 μM of **181** suppressed the proliferation of MCF7 cells (Fig. 4h), while not affecting the ERα negative MDA-MD-231 cells (Fig. 4i).

### Proximity Ligation Assay visualizes the stabilized ERα/14-3-3 PPI, *in situ*

To visualize the direct interaction, as also observed via the nanoBRET (Fig. 4f), between endogenous ERα and 14-3-3 *in situ*, we next performed Proximity Ligation Assays (PLA) in MCF-7 cells (Fig. 5a, Fig. S16-S21). Using PLA, PPIs can be detected by using primary antibodies for each protein followed by conjugation of single-stranded oligonucleotides. When the two proteins are in close proximity (at distances < 40 nm), the oligonucleotides can hybridize followed by PCR amplification and addition of fluorescent probes, allowing the visualization of the spots of proximity by fluorescence microscopy.^35^ MCF-7 cells were treated for 24 hours with 30 μM of FC-A or FC-NAc, or with 1 μM of the synthetic compounds **85, 163** and **181**, followed by fixation and PLA sample preparation for detecting the ERα/14-3-3 PPI. Image analysis for PLA puncta recognition showed an increase in PLA puncta counts in the presence of each stabilizer, compared to the vehicle control (Fig. 5a). This result suggests that all stabilizers enhance the interaction between ERα and 14-3-3, and most prominently in the cytoplasm of the MCF-7 cell. Remarkably, the biochemically very weakly active compound **85** also showed an increase in PLA puncta, which may indicate that PLA is a highly sensitive detection method.

**Fig. 5.**
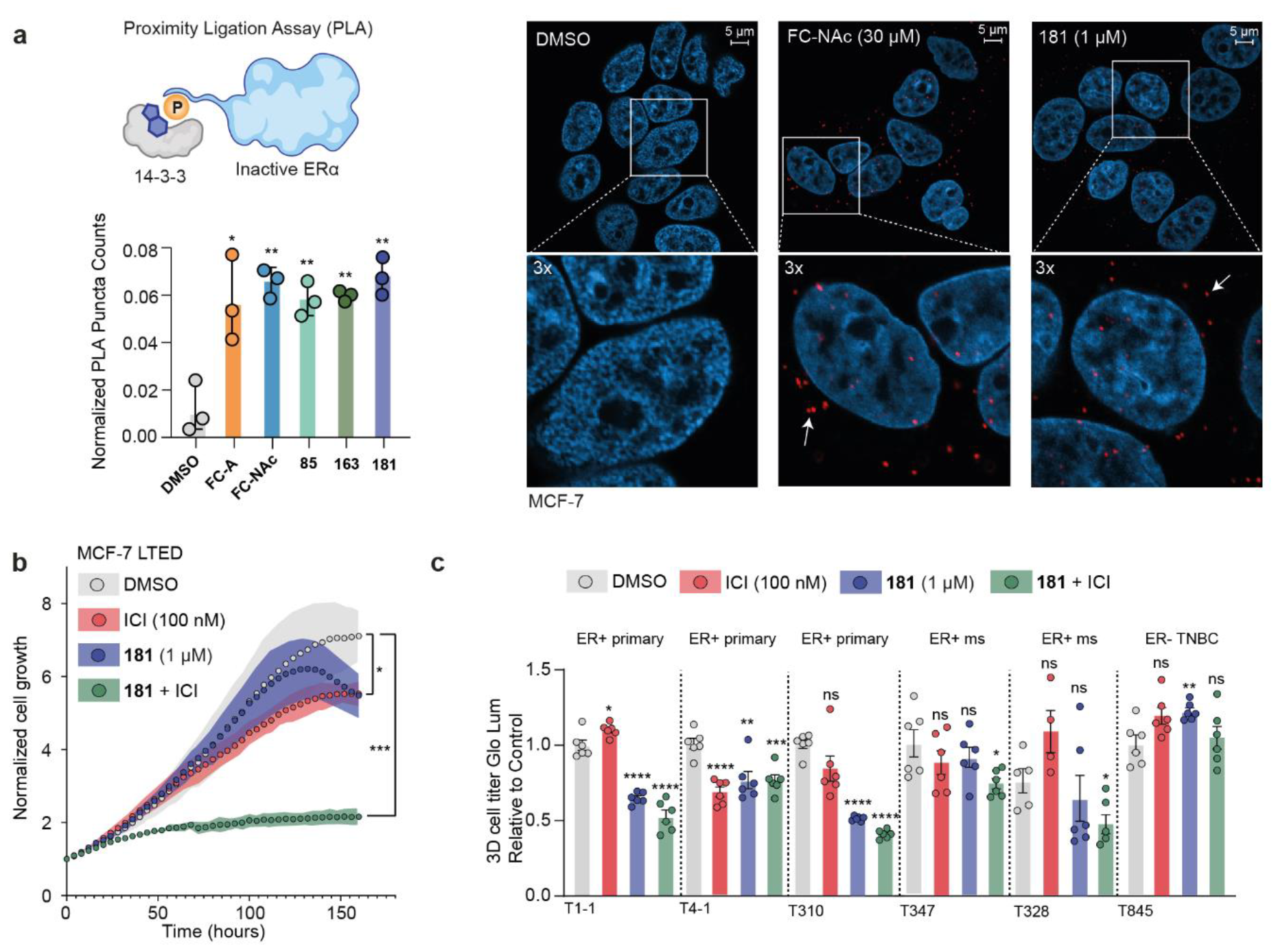
Compound 181, in combination with ICI treatment, suppresses endocrine-resistant cell and organoid proliferation. **a**, Duolink proximity ligation assay (PLA) for protein interactions between ERα and (pan) 14-3-3 in breast cancer cell line MCF-7, when treated with vehicle control DMSO, FC-A (30 μM), FC-NAc (30 μM), **85** (1 μM), **163** (1 μM) or **181** (1 μM). Left: bar graph of PLA puncta counts (mean ± SD, n=3). Right: examples of images of MCF-7 cells treated with DMSO, FC-NAc or **181**. Each red spot represents a single interaction (see arrow for example) and DNA was stained with DAPI (blue). For all images see Fig. S15-S20. **b**, Cellular proliferation assay in MCF-7 LTED cells with DMSO (grey), ICI (100 nM, red), **181** (1 μM, purple) or combination treatment of ICI (100 nM) and **181** (1 μM) (green). (mean ± SD, n = 3 biologically independent, with each 2 technical replicates). **c**, Bar charts of organoid culture proliferation subjected to vehicle control DMSO (grey), ICI (100 nM, red), **181** (1 μM), or combination of ICI (100 nM) and **181** (1 μM) (green). (mean ± SD, n = 4-6, biologically independent). ms = metastatic, TNBC = triple negative breast cancer. (unpaired t-test: *p≤0.05, **p≤0.005 ***p≤0.001, ****p≤0.0001).

### Compound 181 suppresses ERα activity in an endocrine resistant setting

The most potent synthetic stabilizer **181** was further tested for inhibiting cancer-cell growth in an endocrine resistant setting. Cell proliferation experiments were performed on Long-Term Estrogen Deprived (LTED) MCF-7 cells (Fig. 5b) that proliferate independently of estrogens while being dependent on ERα expression. Such cells model resistance to aromatase inhibitor treatment, a phenotype often found in MBC patients.^36^ Compound **181** (1 μM) and ICI (100 nM) individually weakly suppressed MCF-7 LTED proliferation, while the combination of **181** and ICI at these doses resulted in a very impressive, near complete, anti-proliferative effect (Fig. 5b). Further, we analyzed the effect of **181** on cell proliferation of ERα Y537S and ERα D538G cells, showing a 50% reduction of growth at ∼4 μM of **181** (Fig. S21).

Finally, to exemplify the translational potential of our findings, we leveraged organoids derived from patient tumors.^37^ The organoid models included treatment-naïve, endocrine therapy-resistant forms, and a Triple Negative Breast Cancer (TNBC) model reflecting the clinical diversity (Fig. 5c, Fig. S23). The organoids were subjected to the fusicoccanes (FC-A and FC-NAc, both 30 μM), and to the chloro-acetamides (**85, 163**, or **181**, all 1 μM), with and without ICI (100 nM) (Fig. S23). Neither the treatment naïve nor resistant organoid cultures responded to ICI alone, except for line T4-1 (Fig. 5c). In contrast, **181** induced growth inhibition in all treatment naïve organoid cultures. Interestingly, **181** in combination with ICI treatment, suppressed the proliferation of all organoid lines, except for the ERα negative organoid culture T845, derived from TNBC, serving as a negative control.

## Discussion

The major treatment modality for hormonal breast cancer involves blocking ERα activity, typically accomplished by endocrine treatment, including Aromatase Inhibitors (AIs) or by targeting the pocket buried within the LBD of the ERα protein using SERMs (e.g., tamoxifen) or SERDs (e.g., fulvestrant/ICI).^3^ Notwithstanding, drug resistance often arises and one example is due to gene mutations in this binding pocket.^4^ Therefore, attention has been shifted towards the identification of alternative small-molecule modulators that target ERα outside this ligand binding pocket, like the coregulator binding groove^38^ or the ERα/DNA binding interface.^39,40^ Novel LBD-targeting drugs can also be advantageous as demonstrated by the ERα CRBN-based PROTAC (ARV-471)^41^ and covalent cysteine-530 targeting ERα antagonist (HB-6545)^42^, both being currently evaluated in clinical trials (NCT05654623, NCT03250676, NCT04568902, NCT04288089).

Here we focused on targeting a composite pocket created by binding of the suppressive 14-3-3 protein to the C-terminal F-domain of ERα, using molecular glues. We employed an integrated approach to investigate and correlate the biochemical and biological consequences of ERα/14-3-3 PPI stabilization by studying three fusicoccin natural product derivatives with a range of stabilization properties^24^ and fully synthetic covalent stabilizers with low micromolar potency.^25^ We showed that the enhanced semi-synthetic fusicoccin derivative, FC-NAc, is the most potent FC-derived stabilizer for the ERα/14-3-3 PPI reported so far. FC-NAc featured superior suppression of both transcriptional and proliferative effects of ERα action, compared to the original fusicoccin-A (FC-A). RNA-sequencing revealed that both FC-A and FC-NAc indeed suppressed expression of E2/ERα early and late genes, while the weak-stabilizer FC-NAg was inactive.

Because the chemical structure of fusicoccanes impedes straightforward medicinal chemistry optimization, we also tested the biological effect of synthetic covalent stabilizers for the ERα/14-3-3σ PPI. We demonstrated that the most potent stabilizer, compound **181**, suppressed the transcriptional activity of ERα at significantly lower concentrations than the fusicoccanes (1 μM versus 30 μM, respectively), resulting in an inhibition of MCF-7 cell proliferation and an increase in Thr594 ERα phosphorylation, associated with an enhanced ERα/14-3-3 PPI formation. Interestingly, these synthetic stabilizers bind covalently to Cys38 in the 14-3-3 pocket, which is only present in the sigma (σ) isoform of 14-3-3. Multiple studies found a downregulation or degradation of 14-3-3σ expression in some breast cancers, which often related to poor prognosis, and 14-3-3σ was therefore considered to be a tumor suppressor.^43–51^ However, 14-3-3σ is still commonly expressed in ERα-positive breast cancer (as shown in the Human Protein Atlas) and the expression level of ERα is low as compared to 14-3-3σ, thus providing value for exploring molecular glues that can shift the ERα/14-3-3 binding equilibrium towards the sigma isoform of 14-3-3, thereby enforcing its tumor suppressive role.^52^

The most common ERα gene mutations linked to endocrine drug resistance are Y537S and D538G.^7^ The distant location of these mutations compared to the ERα/14-3-3 PPI pocket targeted by our stabilizers would potentially enable the PPI stabilization mechanism to be active even in these point-mutated cell lines. Indeed, we demonstrated that both FC-A and FC-NAc suppressed mutant ERα transcriptional activity, which could be further enhanced by addition of ICI. The FCs and **181** reduced the proliferation of Y537S and D538G ERα mutant cell lines, particularly in combination with ICI. **181** was also effective in Long Term Estrogen Deprived (LTED) MCF7 cells and in endocrine-resistant patient-derived organoids, either alone or in combination with ICI, dependent on the organoid mutational background. It is noteworthy that our compounds do show some reduced potency in the ERα-altered lines; given the physical separation of the LBD and the C-terminal 14-3-3 binding site, why are the molecular glues less active? Y537S mutated ERα has been shown to result in a modified transcriptional complex, with an increased association of FOXA1 and GREB1.^36^ Thus, the enhanced formation of modified transcriptional complexes of the mutated ERα constructs, potentially in combination with alternative cellular localization or steric hindrance by altered interacting proteins, may counteract 14-3-3 binding. Nevertheless, we show that small-molecule stabilization can shift the equilibrium to formation of ERα/14-3-3 complexes, which can be exploited to target endocrine resistant breast cancer, perhaps in combination with endocrine antagonists like ICI.

Finally, proximity ligation assays (PLA), in line with nanoBRET data, showed an increase in PLA puncta of the ERα/14-3-3 PPI upon addition of both natural and synthetic stabilizers. These PLA puncta were both visible in the cytoplasm and nucleus of MCF-7 cells. ERα is predominantly found in the nucleus, both in hormone stimulated and untreated cells.^53^ However, while the steady state distribution of ERα is heavily nuclear enriched, ERα does shuttle between the nucleus and cytosol.^54^ It is known that 14-3-3 proteins can promote either the cytoplasmic-, or nuclear localization of its binding partners^55^, and these data suggest that 14-3-3 can suppress ERα activity by interacting in the cytoplasm of breast cancer cells. Clearly, additional studies involving the cellular localization of ERα upon stabilization to 14-3-3 will be required to further validate this intriguing observation.

Taken together, we have learned that the suppressive role of 14-3-3 on ERα activity in breast cancer proliferation can be enhanced by small molecule molecular glues of this PPI, as illustrated using two different classes of compounds – natural product analogs and 14-3-3σ-specific covalent synthetic molecules. For both chemotypes a clear Structure-Activity-Relationship could be shown ranging from biochemical and structural data to cellular systems of diverse complexity, including breast cancer organoids. Importantly, this unique mechanism of action could also be exploited in endocrine resistant conditions across the two diverse chemotypes. This methodology of PPI stabilization can not only be applied to the ERα/14-3-3 PPI but may be translated to other nuclear receptors as well. For example, the Androgen Receptor (AR) has been reported to bind to 14-3-3 via its N-terminal domain (NTD)^56^, potentially offering an orthogonal strategy of targeting castration-resistant prostate cancer (CRPC).^57^ Here we demonstrate the foundations for the development of a novel class of ERα-targeting compounds - validated by natural products and the first cellularly active covalent ERα/14-3-3σ stabilizers - providing a first proof-of-concept for their development towards potential future clinical applications.

## Acknowledgments

We would like to thank Yusuke Higuchi for his kind gift of FC-A, deacetylated FC-A and the aglycon derivative; FC-NAg. We acknowledge Sebastian Andrei for the semi-synthesis of some of the fusicoccanes used in this manuscript. We thank the NKI genomics core facility for technical assistance with the RNA sequencing data and Sebastian Gregoricchio is acknowledged uploading RNA sequencing data to the Geo database. Bas Rosier is kindly thanked for useful discussions and insights in story conceptualization. We acknowledge DESY (Hamburg, Germany), a member of the Helmholtz Association HGF, for the provision of experimental facilities. Parts of this research were carried out at PETRA III and we would like to thank Anja Burkhardt and Sofiane Saouane for their assistance in using beam P11. Beamtime was allocated for proposals 11006723 and 11009075. The research described was funded by the Netherlands Organization for Scientific Research (NWO) through Gravity program 024.001.035 and ECHO grant 711.018.003.

## Author contributions

E.J.V. conceived the work, designed and performed experiments, and analyzed the data with contributions from M.D.C, B.A.S., M.R.A., C.O., W.Z. and L.B. RNA-sequencing data was processed and analyzed by J.S. The chloroacetamide containing compounds were synthesized and verified by M.K. The NanoBRET assay was performed and analyzed by J.M.V. The proximity ligation assay was performed and analyzed by S.N.P. and J.B.B., with contributions from O.C.M. The organoid experiments were performed by L.L, L.Y., and D.V. Specific MCFR-7 cell lines were prepared by L.B. and S.A. The synthesis of FC-NAc was performed by P.C. and G.M. E.J.V. wrote the manuscript with contributions from M.R.A., C.O., W.Z., and L.B.

## Supporting information

Supplementary figures and tables, extended data.

## Methods

### Protein expression and purification

The 14-3-3*σ* isoform with a truncated C-terminus after T234 (ΔC: to enhance crystallization) and a N-terminal His6-tag was expressed in NiCo21 (DE3) competent E.coli (new England biolabs Inc.) from a pPROEX HTb expression vector. After transformation following manufacturer’s instructions, single colonies were picked to inoculate 30 mL precultures (LB), which were added to 1.5 L 2XYT medium after overnight growth at 37 °C, 250 rpm. Expression was induced upon reaching OD6000.5-0.6 by adding 400 μM IPTG. After overnight expression at 18 °C, 140 rpm, cells were harvested by centrifugation at 8000 rpmn and resuspended in lysis buffer (50 mM Tris, pH 8.0, 300 mM NaCl, 10 mM imidazole, 5 mM MgCl2, 1 mM PMSF, 250 μM TCEP). The His6-tagged proteins were first purified by Ni-affinity chromatography (HisTrap HP column, GE) (Elution buffer 50 mM Tris, pH 8.0, 300 mM NaCl, 250 mM imidazole, 250 μM TCEP), followed by His-tag cleavage by TEV protease during dialysis (25 mM HEPES pH 7.5, 200 mM NaCl, 5% glycerol, 10 mM MgCl2, 250 μM TCEP overnight at 4 °C. The flowthrough of a second HisTrap column was subjected to final purification step by size-exclusion chromatography (Superdex 75, GE) (SEC buffer 25 mM HEPES pH 7.5, 100 mM NaCl, 10 mM MgCl2, 250 μM TCEP). The protein was concentrated to ∼60 mg/mL, analyzed for purity by SDS-page and Q-Tof LC/MS and aliquots flash-frozen for storage at -80 °C.

### Peptides

ERalpha-peptide for X-ray crystallography and FAM-labeled peptides were ordered from GenScript Biotech Corp. Sequence of ERalpha-peptide: Ac- or 5-FAM-AEGFPA{pT}V-COOH (8mer).

### Compounds

FC-A, deacetylated FC-A and FC-NAg was a kind gift from Yusuke Higuchi. All other fusiccocin-derivatives were synthesized as described previously by Andrei *et al*.^58^ The chloro-acetamides were synthesized as described by Konstantinidou *et al*.^25^ Fulvestrant (ICI), Estradiol (E2) and MG-132 were purchased from MedChemExpress (HY-13636, HY-B0141, HY-13259, respectively).

### Fluorescence Anisotropy

Fluorescein-labeled peptides (5-FAM), 14-3-3σ FL protein, the fusicoccanes (10 mM stock solution in DMSO) were diluted in buffer (10 mM HEPES, pH 7.5, 150 mM NaCl, 0.1% Tween20, 1 mg/mL Bovine Serum Albumin (BSA; Sigma-Aldrich). Final DMSO in the assay was always 1%. Dilution series of 14-3-3 proteins or compounds were made in black, round-bottom 384-microwell plates (Corning) in a final sample volume of 10 μL in triplicates.

Compound titrations were made by titrating the compound in a 2-fold dilution series (starting at 100 μM) to a mix of fluorescein-labelled peptide (10 nM) and 14-3-3σ (concentration at EC_20_ value of the protein-peptide complex; 1 μM for ERα). Fluorescence anisotropy measurements were performed directly.

Protein titrations were made by titrating 14-3-3σ in a 2-fold dilution series (starting at 300 μM) to a mix of fluorescein-labelled peptide (10 nM) and DMSO or compound (100 μM). Fluorescence anisotropy measurements were performed directly.

Protein 2D titrations were made by titrating 14-3-3σ in a 2-fold dilution series (starting at 250 μM) to a mix of fluorescein-labelled peptide (10 nM) against varying fixed concentrations of compound (2-fold dilution, starting at 250 μM), or DMSO. Fluorescence anisotropy measurements were performed directly.

Fluorescence anisotropy values were measured using a Tecan Infinite F500 plate reader (filter set lex: 485 ± 20 nm, lem: 535 ± 25 nm; mirror: Dichroic 510; flashes:20; integration time: 50 ms; settle time: 0 ms; gain: 55; and Z-position: calculated from well). Wells containing only FAM-peptide were used to set as G-factor at 35 mP. Data reported are at endpoint. EC_50_ and apparent K_d_ values were obtained from fitting the data with a four-parameter logistic model (4PL) in GraphPad Prism 7 for Windows. Data was obtained and averaged based on either three (compound titrations) or two (protein titrations) independent experiments.

### X-ray Crystallography; data collection and refinement

14-3-3 protein (470 μM, 12.5 mg/mL) was mixed with ERα-peptide (1:2 molar stoichiometry; 940 μM) and incubated in crystallization buffer (20 mM HEPES pH 7.4, 2 mM MgCl2, 2 mM BME) for a few minutes at RT before setting up for sitting drop crystallization in MRC crystallization plates (Swissci) with a custom crystallization liquor-grid (0.095 M HEPES (pH 7.1, 7.3, 7.5, 7.7), 0,19 M CaCl2, 5% glycerol, 24-29% PEG 400). Crystals grew at 4 °C within 1 week. Soaking of crystals was performed by mixing 0.4 μL FC-derivatives from 10 mM stock solutions in DMSO in 3.6 μL mother liquor, which was then added to crystal-containing drops. Soaked crystals were fished after overnight incubation and flash-frozen in liquid nitrogen. Diffraction data were collected at the Deutsche Electronen Synchrotron (DESY, PETRA-III beamline), equipped with a Dectris Pilatus 6M-f detector. Data was processed using the CCP4i2 suite (version 8.0.003)^59^. Data integration was done using Xia2Dials^60^. The data was phased with MolRep^61^, using 4JC3 as a template. Presence of soaked ligands was verified by visual inspection of the Fo-Fc and 2Fo-Fc electron density maps in COOT (version 0.9.6)^62^. eLBOW^63^ was used to generate the structures and restraints of the soaked ligands, followed by model rebuilding and refinement using phenix.refine^64,65^ from the Phenix software suite (version 1.19.2-4158) and Coot. The images were created using the PyMol Molecular Graphic System (Schrödinger LLC, version 2.2.3). The structures were deposited in the protein data bank (PDB) with IDs: 8BZD (FC-dAc), 8BZE (FC-J), 8BZF (FC-J acetonide), 8BZG (FC-31), 8C0L (FC-THF), 8BZH (FC-NAc) and 8BZT (FC-NAg).

### Cell lines

MCF-7 (Michigan Cancer Foundation-7) human breast carcinoma cell line was obtained from ATCC. MCF-7 Y537S (clone A4, 1 and 8) and D538G (clone 5.1, 3, and 4) mutant cell lines were generated using CRISRP-Cas9, as previously described.^32^ T47D, ZR-75-01, MDA-MB-231 and HEK293T cells were purchased from ATCC. Cell lines were maintained in regular Dulbecco’s modified Eagle’s medium (DMEM) supplied with 10% fetal bovine serum (FBS) and 1% penicillin/streptomycin, and cultured at 37 °C and with 5% CO_2_. Three days before all experiments requiring an E2 induction, cells were placed in hormone-depleted media. Hormone-depleted media consisted of phenol-red free DMEM (Thermo Fisher Scientific), 5% dextran and charcoal-stripped fetal bovine serum, 1% penicillin-streptomycin and 1% L-glutamate. All cell lines were tested negative for mycoplasma contamination.

### Cell Titer Blue (CTB) viability assay

Hormone-deprived MCF-7 and MDA-MB-231 cells were seeded at 2,500 cells per well in a 96-well plate. Cells were pretreated 1h with the indicated concentration of fusicoccanes or chloro-acetamides (starting at a concentration of 50 μM, dilution 1:3) or ICI (starting at a concentration of 1 μM, dilution 1:3). After 1 hour of incubation, 10 nM of E2 was added to the wells containing the MCF-7 cells to induce proliferation. After one week of incubation, cell viability was measured by addition of resazurin according to manufacturer’s instructions (Promega).

### Luciferase Reporter Assay

Hormone-deprived MCF-7 cells were seeded at 10,000 cells per well in a 96-well plate. Cells were transfected for 6 hours with 80 ng 3xERRE-ERE-luciferase containing codon-modified firefly luciferase (Addgene #37852) and 16 ng pRL-SV40P (Addgene #27163) per well using lipofectamine 3000 (Thermo Fisher), followed by an overnight treatment with the indicated concentration of fusicoccanes, chloroacetamides or ICI. The following day the cells were stimulated with the indicated concentration of E2 for 24 h, after which the ERα activity was determined with a Dual-Luciferase Reporter Assay (Promega), according to the manufacturer’s instructions. The ERE-luciferase signal was first normalized over the Renilla-luciferase signal.

### Western blotting

Cells were seeded in a 6-well plate (150.000 cells/well) in full medium. The next day the cells were treated with the fusicoccanes (30 μM) or **181** (1 μM) for 72 hours. Fulvestrant (ICI, 100 nM) and/or MG-132 (5 μM) was added one day before cell harvesting. Cells were then washed once with PBS and harvested in Laemmli buffer (60 mM Tris pH 6.8, 10% v/v glycerol, 2% SDS) supplemented with protease inhibitors (cOmplete, EDTA-free Protease Inhibitor Cocktail, Roche) and phosphatase inhibitors (100 μM Na_3_VO_4_ CAS:13721-39-6 ThermoFisher, 100 mM NaF Jena Bioscience), followed by sonication (Bioruptor® Pico Diagenode). Total protein content was quantified by BCA assay (Pierce BCA Protein Assay Kit, ThermoFisher). Twenty micrograms of reduced protein per lane was assayed on 10% polyacrylamide gels and transferred to a PVDF membrane according to manufacturer’s specifications. The PVDF membrane was blocked for 1 h with 5% BSA in TBST buffer containing 0.1% Tween 20 followed by overnight incubation at 4 °C with primary antibody for: ERα (MA5-14104, dilution 1:400, Thermo Fisher Scientific) and PR (sc-7208, dilution 1:1000, santa-cruz). The antibody raised against the phosphorylated Thr594 of ERα was prepared by GL Biochem as described previously (dilution 1:100).^13^ Actin served as loading control (ab6276, dilution 1:5000, abcam). PVDF membranes were washed 3x over a period of 30 min with TBST before incubating the membranes with anti-mouse or anti-rabbit fluorescently labeled secondary antibody (1:10,000, IRDye®) for 1 h at room temperature, followed by 3x washes with TBST over a period of 30 min. Blots were developed on a LI-COR Odyssey imaging system according to manufacturer’s specification and analyzed using Image Studio Lite (version 5.2.5).

### RNA-sequencing

300.000 MCF-7 WT cells were plated in 6-well dishes and grown in hormone-depleted media for 6 days containing 30 μM FC-A, FC-NAc, FC-NAg, or DMSO as vehicle control, followed by E2 stimulation (10 nM) for 6 hours. Cells were washed with phosphate-buffered saline (PBS; Thermo Fisher Scientific) and treated with RTL plus (Qiagen) containing 1% beta-mercaptoethanol (Sigma-Aldrich) for cell lysis. Cells were passed through a 21-gauge needle and syringe (Sigma-Aldrich) several times to facilitate lysis of genomic DNA. RNA was then extracted using a Quick-RNA Miniprep kit (Zymo Research). RNA was quantified and poly(A) selected RNA-seq libraries were generated using a KAPA Stranded mRNA-seq kit (KAPA Biosystems) with 500 ng RNA per sample as starting material. Libraries were sequenced on the Illumina HiSeq 2500 platform. Resulting RNA-seq data were quantified using salmon v1.4 in mapping-based mode against a decoy-aware indexed GRCh38 transcriptome using Gencode v36 transcript annotations with default k-mer length and –validateMApping flag. The resulting quantification files were imported into R v4.1, summarized to gene-level with tximport, and differential expression analysis was performed with DESeq2. Gene Set Enrichment Analysis (GSEA) was performed using the enricher function from clusterProfiler against msigdbr Hallmarks gene set with the differentially expressed genes ranked by log2foldChange * adjusted p-value. Only significant pathways with a benjamini Hochberg adjusted p-value less than 0.05 were plotted, and activated pathways indicate a positive log2foldChange while suppressed pathways indicate a negative log2foldchange. RNA-seq data is submitted to GEO and was allocated submission number: GSE227960

### Proliferation assay

Hormone-deprived MCF-7 cells (ESR1 WT, Y537S, D538G mutants, LTED), T47D, ZR-75-01 and MDA-MB-231 cells were seeded at 2,500 cells per well in a 96-well plate. Cells were pretreated 1h with the indicated concentration of fusicoccanes (FC), chloroacetamides or DMSO as control, before the addition of 10 nM E2. Cell proliferation was measured kinetically every 4 hours as an increase in cell confluence. Cell confluence was determined by analysis of phase-contrast images, using the IncuCyte FLR (Essen BioScience). The algorithm used to analyze images for cell confluence was from the IncuCyte software, build 1001A Rev2.

### NanoBRET assay

NanoBRET assays were performed as described by Promega. HEK293T cells were cultured as described above and transfected with a 1:10 ratio of Nluc-ERa:14-3-3σ-HaloTag plasmid for 48 hours using jetOPTIMUS transfection reagent (Polyplus). The media was replaced to Gibco FluoroBrite DMEM (phenol red-free) with 4% FBS and 1% penicillin/streptomycin. Cells were then seeded at 8,000 cells per well in a 384-well plate (Corning #3570) and treated with 100 nM HaloTag NanoBRET 618 Ligand (Promega) or equivalent volume of DMSO as a no acceptor negative control. Following plating, cells were treated for 24 hours with compound in 1:2 dilution series starting at 20 μM chloro-acetamides (0.35% DMSO final concentration). After 24 hours, the BRET signal was read using an EnVision XCite 2105 plate reader at 618 nm (HaloTag) and 460 nm (Nluc). The final corrected NanoBRET ratio was calculated using the following equation:

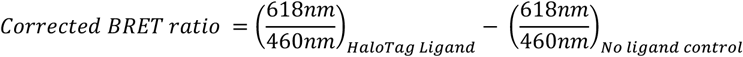

The BRET ratios were normalized to samples treated with DMSO instead of the chloro-acetamide compounds.

### Proximity Ligation Assay (PLA)

Hormone deprived MCF-7 cells were seeded in 24-well plates (100.000 cells/well), followed by stabilizer treatment the next day (30 μM of FC-A or FC-NAc; 1 μM for 85, 163 or 181). After 24 hours of treatment, the cells were washed and fixated with 4% paraformaldehyde. In situ PLA was performed using Duolink PLA reagents (Sigma Aldrich) largely according to the manufacturer’s instructions. For experiments on MCF-7 cell cultures, samples were incubated in Duolink blocking buffer at 37 °C for 1 h. We then applied primary antibodies, (rabbit anti-Estrogen Receptor alpha antibody [ab3575] at a 1:2000 dilution and mouse pan 14-3-3 Antibody (B-8) [sc-133233] at a 1:100 dilution, in antibody diluent (0.3% triton X-100 and 0.1% PBS solution) overnight at 4 °C. After washing 3 × 10 min in PLA wash buffer A, we probed the primary antibodies using Duolink PLA probes rabbit PLUS and mouse MINUS at 37 °C for 1 h and then used Duolink In Situ Detection Reagents Red (Sigma Aldrich) to detect the proximity between the proteins of interest. To image PLA signals a total of 10 images were acquired per coverslip (3 coverslips per condition) with the confocal LSM 900 Airyscan (Zeiss), using the 63X objective magnification with immersion oil. Images were processed in Fiji^66^. Images were scaled to 1892 × 1892 and converted to 8-bit. The DAPI channel was binarized using a inclusion threshold of 20 – 255. Sub-threshold ‘dark spots’ inside of nuclei were accounted for with the “Fill holes” function. The PLA channel was binarized using a percentile threshold, set at the upper 0.5% percentile of pixel intensities. The binarized PLA layer was de-noised using “Despeckle” and “Smooth”. The layer was then re-binarized, followed by “Despeckle” and “Watershed”. Using the binarized DAPI and PLA layers, puncta inside and outside the nuclei were inferred using the “AND” and “Subtract” functions, respectively. Finally, a stack was created containing the binarized DAPI layer, the binarized PLA spots inside and outside the nuclei, and the raw PLA layer. Measurements of the stack were taken using “Analyze Particles” with arguments “size=0.01-Infinity circularity=0-1.00 display clear overlay add stack”. Measurement options included “Area”, “Standard Deviation”, “Mean gray value”, “Min & max gray value”, and “Shape descriptors”.

Measurements were then imported and processed in R version 4.3.1 (R Core Team, 2023). The number of nuclei and the sum of nucleic area was recorded per image. The measured PLA puncta were filtered on area and shape parameters. Thresholds used were 0.1 < Area < 0.35, Circ. > 0.85, AR < 1.75, 0.88 < Solidity < 0.95. Thereafter, puncta were filtered on the max gray value recorded in the raw PLA layer. Puncta with a raw max gray value higher than 50 were retained. Puncta were tallied per image, both inside and outside the nuclei. Total puncta per image was obtained by addition of the inside and outside counts. Finally, the total counts per image were normalized against the sum of nucleic area.

### Organoids

Organoid experiments were conducted at RCSI University of Medicine and Health Sciences. We developed tumour-derived organoids following ethical guidelines approved by RCSI’s Institutional Review Board. Our approach involved standard organoid techniques for creating lines from tumour samples, adding estradiol for ER+ tumours (PMID: 2922478033208158). For intervention experiments, we dissociated mature organoids and cultured them in organoid-specific media, incorporating 5% Cultrex® Reduced Growth Factor Basement Membrane Matrix (BME, Trevigen, 3533-001-02). Organoids underwent treatment with either vehicle or specified, compounds at set concentrations (N=6; biological replicates). Cell viability was assessed 7 days post-treatment using the CellTiter-Glo® 3D Cell Viability assay (Promega).

## Data availability

Crystal structures described in this manuscript have been deposited to the PDB with following IDs: 8BZD, 8BZE, 8BZF, 8BZG, 8C0L, 8BZH, 8BZT. RNA-sequencing data is submitted to GEO and was allocated submission number: GSE227960.

## Code availability

Algorithms used by custom analysis code for RNA sequencing and PLA data analysis are described in detail in the Methods. Code is available upon reasonable request.

## References

1. Siegel, R. L. Cancer statistics, 2022. 72, 7–33 (2022).

2. Liu, Y., Ma, H. & Yao, J. ERα, a key target for cancer therapy: A review. Onco. Targets. Ther. 13, 2183–2191 (2020).

3. Aggelis, V. & Johnston, S. R. D. Advances in Endocrine - Based Therapies for Estrogen Receptor - Positive Metastatic Breast Cancer. Drugs (2019).

4. Davies, C. et al. 20-Year Risks of Breast-Cancer Recurrence after Stopping Endocrine Therapy at 5 Years. N. Engl. J. Med. 377, 1836–46 (2017).

5. Saatci, O., Huynh-Dam, K. T. & Sahin, O. Endocrine resistance in breast cancer: from molecular mechanisms to therapeutic strategies. J. Mol. Med. 99, 1691–1710 (2021).

6. Hanker, A. B., Sudhan, D. R. & Arteaga, C. L. Overcoming Endocrine Resistance in Breast Cancer. Cancer Cell 37, 496–513 (2020).

7. Fanning, S. W. et al. Estrogen receptor alpha somatic mutations Y537S and D538G confer breast cancer endocrine resistance by stabilizing the activating function-2 binding conformation. Elife 5, 1–25 (2016).

8. Jeselsohn, R., Buchwalter, G., Angelis, C. De, Brown, M. & Schiff, R. ESR1 mutations as a mechanism for acquired endocrine resistance in breast cancer. Nat. Rev. Clin. Oncol. 12, 573–583 (2015).

9. Arnesen, S. et al. Estrogen Receptor Alpha Mutations in Breast Cancer Cells Cause Gene Expression Changes through Constant Activity and Secondary Effects. Cancer Res. 81, 539–551 (2021).

10. Herzog, S. K. & Fuqua, S. A. W. ESR1 mutations and therapeutic resistance in metastatic breast cancer: progress and remaining challenges. Br. J. Cancer 126, 174–186 (2022).

11. Padrão, N. A., Mayayo-Peralta, I. & Zwart, W. Targeting mutated estrogen receptor alpha: Rediscovering old and identifying new therapeutic strategies in metastatic breast cancer treatment. Curr. Opin. Endocr. Metab. Res. 15, 43–48 (2020).

12. Morrison, D. K. The 14-3-3 proteins: integrators of diverse signaling cues that impact cell fate and cancer development. Trends Cell Biol. 19, 16–23 (2009).

13. De Vries-van Leeuwen, I. J. et al. Interaction of 14-3-3 proteins with the Estrogen Receptor Alpha F domain provides a drug target interface. Proc. Natl. Acad. Sci. 110, 8894–8899 (2013).

14. Mohammed, H. et al. Endogenous purification reveals GREB1 as a key Estrogen Receptor regulatory factor. Cell Rep. 3, 1–16 (2013).

15. Mohammed, H. et al. Progesterone receptor modulates ER a action in breast cancer. Nature 523, 313–319 (2015).

16. Campbell, T. M., Castro, M. A. A., Oliveira, K. G. De, Bruce, A. J. & Meyer, K. B. ER alpha binding by transcription factors NFIB and YBX1 enables FGFR2 signalling to modulate estrogen responsiveness in breast cancer. Cancer Res. 78, 410–421 (2018).

17. Papachristou, E. K. et al. A quantitative mass spectrometry-based approach to monitor the dynamics of endogenous chromatin-associated protein complexes. Nat. Commun. 9, 1–13 (2018).

18. Schreiber, S. L. The Rise of Molecular Glues. Cell 184, 3–9 (2021).

19. Gerry, C. J. & Schreiber, S. L. Unifying principles of bifunctional, proximity-inducing small molecules. Nat. Chem. Biol. 16, 369–378 (2020).

20. Schapira, M., Calabrese, M. F., Bullock, A. N. & Crews, C. M. Targeted protein degradation: expanding the toolbox. Nat. Rev. Drug Discov. 18, 949–963 (2019).

21. Yamshon, S. & Ruan, J. IMiDs New and Old. Curr. Hematol. Malig. Rep. 14, 414–425 (2019).

22. Sun, X. et al. Protacs: Great opportunities for academia and industry. Signal Transduct. Target. Ther. 4, (2019).

23. Chen, B. et al. A Two-Phase Approach to Fusicoccane Synthesis To Uncover a Compound That Reduces Tumourigenesis in Pancreatic Cancer Cells. Angew. Chemie - Int. Ed. 61, 1–8 (2022).

24. Marra, M., Camoni, L., Visconti, S., Fiorillo, A. & Evidente, A. The surprising story of fusicoccin: A wilt-inducing phytotoxin, a tool in plant physiology and a 14-3-3-targeted drug. Biomolecules 11, 1–29 (2021).

25. Konstantinidou, M. et al. Structure-based Optimization of Covalent, Small-Molecule Stabilizers of the 14-3-3σ/ERα Protein-Protein Interaction from Nonselective Fragments. J. amer (2023). doi:10.1021/jacs.3c05161

26. Barrow, K. D., Barton, D. H. R., Chain, E., Bageenda-Kasujja, D. & Mellows, G. Fusicoccin. Part IV. The structure of fusicoccin J. J. Chem. Soc. Perkin Trans. 1 877–883 (1975).

27. De Vink, P. J. et al. Cooperativity basis for small-molecule stabilization of protein-protein interactions. Chem. Sci. 10, 2869–2874 (2019).

28. Anders, C. et al. A semisynthetic fusicoccane stabilizes a protein-protein interaction and enhances the expression of K+ channels at the cell surface. Chem. Biol. 20, 583–593 (2013).

29. Andrei, S. A. et al. Rationally Designed Semisynthetic Natural Product Analogues for Stabilization of 14-3-3 Protein–Protein Interactions. Angew. Chemie Int. Ed. 57, 13470–13474 (2018).

30. Petz, L. N., Ziegler, Y. S., Loven, M. A. & Nardulli, A. M. Estrogen receptor α and activating protein-1 mediate estrogen responsiveness of the progesterone receptor gene in MCF-7 breast cancer cells. Endocrinology 143, 4583–4591 (2002).

31. Dustin, D., Gu, G. & Fuqua, S. A. W. ESR1 mutations in breast cancer. Cancer 125, 3714–3728 (2019).

32. Harrod, A. et al. Genomic modelling of the ESR1 Y537S mutation for evaluating function and new therapeutic approaches for metastatic breast cancer. Oncogene 36, 2286–2296 (2017).

33. Toy, W. et al. Activating ESR1 mutations differentially impact the efficacy of ER antagonists. Cancer Discov 7, 277–287 (2017).

34. Jeselsohn, R. et al. Allele-Specific Chromatin Recruitment and Therapeutic Vulnerabilities of ESR1 Activating Mutations. Cancer Cell 33, 173-186.e5 (2018).

35. Muhammad S Alam. Proximity Ligation Assay (PLA). Curr. Protoc. Immunol. 123, e58 (2018).

36. Martin, L. A. et al. Discovery of naturally occurring ESR1 mutations in breast cancer cell lines modelling endocrine resistance. Nat. Commun. 8, (2017).

37. Sachs, N. et al. A Living Biobank of Breast Cancer Organoids Captures Disease Heterogeneity. Cell 172, 373-386.e10 (2018).

38. Raj, G. V et al. Estrogen receptor coregulator binding modulators (ERXs) effectively target estrogen receptor positive human breast cancers. Elife 6, 1–69 (2017).

39. Moore, T. W., Mayne, C. G. & Katzenellenbogen, J. A. Minireview: Not picking pockets: Nuclear receptor alternate-site modulators (NRAMs). Mol. Endocrinol. 24, 683–695 (2010).

40. Zhang, X. et al. PROTAC Degrader of Estrogen Receptor α Targeting DNA-Binding Domain in Breast Cancer. ACS Pharmacol. Transl. Sci. 5, 1109–1118 (2022).

41. Negi, A., Kesari, K. K. & Voisin-Chiret, A. S. Estrogen Receptor-α Targeting: PROTACs, SNIPERs, Peptide-PROTACs, Antibody Conjugated PROTACs and SNIPERs. Pharmaceutics 14, 2523 (2022).

42. Furman, C. et al. Covalent ERa Antagonist H3B-6545 Demonstrates Encouraging Preclinical Activity in Therapy-Resistant Breast Cancer. Mol. Cancer Ther. 21, 890–902 (2022).

43. Laronga, C., Yang, H. Y., Neal, C. & Lee, M. H. Association of the cyclin-dependent kinases and 14-3-3 sigma negatively regulates cell cycle progression. J. Biol. Chem. 275, 23106–23112 (2000).

44. Hynes, N. E. & Smirnova, T. The 14-3-3σ tumor suppressor has multiple functions in ErbB2-induced breast cancer. Cancer Discov. 2, 19–22 (2012).

45. Ferguson, A. T. et al. High frequency of hypermethylation at the 14-3-3 σ locus leads to gene silencing in breast cancer. Proc. Natl. Acad. Sci. U. S. A. 97, 6049–6054 (2000).

46. Simooka, H., Oyama, T., Sano, T., Horiguchi, J. & Nakajima, T. Immunohistochemical analysis of 14-3-3 sigma and related proteins in hyperplastic and neoplastic breast lesions, with particular reference to early carcinogenensis. Pathol. Int. 54, 595–602 (2004).

47. Moreira, J. M. A., Ohlsson, G., Rank, F. E. & Celis, J. E. Down-regulation of the tumor suppressor protein 14-3-3σ is a sporadic event in cancer of the breast. Mol. Cell. Proteomics 4, 555–569 (2005).

48. Mhawech, P. et al. Downregulation of 14-3-3σ in ovary, prostate and endometrial carcinomas is associated with CpG island methylation. Mod. Pathol. 18, 340–348 (2005).

49. Ye, M. et al. Detection of 14-3-3 sigma (σ) promoter methylation as a noninvasive biomarker using blood samples for breast cancer diagnosis. Oncotarget 8, 9230–9242 (2017).

50. Mei, J. et al. Characterization of the expression and prognostic value of 14-3-3 isoforms in breast cancer. Aging (Albany. NY). 12, 19597–19617 (2020).

51. Huang, Y., Yang, M. & Huang, W. 14-3-3 σ: A potential biomolecule for cancer therapy. Clin. Chim. Acta 511, 50–58 (2020).

52. Liu, Y., Liu, H., Han, B. & Zhang, J. T. Identification of 14-3-3σ as a contributor to drug resistance in human breast cancer cells using functional proteomic analysis. Cancer Res. 66, 3248–3255 (2006).

53. Monje, P., Zanello, S., Holick, M. & Boland, R. Differential cellular localization of estrogen receptor a in uterine and mammary cells. 181, 117–129 (2001).

54. Maruvada, P., Baumann, C. T., Hager, G. L. & Yen, P. M. Dynamic Shuttling and Intranuclear Mobility of Nuclear Hormone Receptors *. J. Biol. Chem. 278, 12425–12432 (2003).

55. Muslin, A. J. & Xing, H. 14-3-3 proteins: regulation of subcellular localization by molecular interference. 12, 703–709 (2000).

56. Ruff, S. E., Vasilyev, N., Nudler, E., Logan, S. K. & Garabedian, M. J. PIM1 phosphorylation of the androgen receptor and 14-3-3 ζ regulates gene transcription in prostate cancer. Commun. Biol. 4, (2021).

57. Monaghan, A. E. & McEwan, I. J. A sting in the tail: The N-terminal domain of the androgen receptor as a drug target. Asian J. Androl. 18, 687–694 (2016).

58. Andrei, S. A. Engineering stabilizers of 14-3-3 protein-protein interactions. (Technical University Eindhoven, 2019).

59. Potterton, L. et al. CCP 4 i 2: The new graphical user interface to the CCP 4 program suite. Acta Crystallogr. Sect. D Struct. Biol. 74, 68–84 (2018).

60. Winter, G. et al. DIALS: Implementation and evaluation of a new integration package. Acta Crystallogr. Sect. D Struct. Biol. 74, 85–97 (2018).

61. Vagin, A. & Teplyakov, A. Molecular replacement with MOLREP. Acta Crystallogr. Sect. D 66, 22–25 (2010).

62. Emsley, P. & Cowtan, K. Coot: Model-building tools for molecular graphics. Acta Crystallogr. Sect. D Biol. Crystallogr. 60, 2126–2132 (2004).

63. Moriarty, N. W., Grosse-Kunstleve, R. W. & Adams, P. D. eLBOW: a tool for ligand coordinate and restraint generation. Acta Crystallogr. Sect. D 65, 1074–1080 (2009).

64. Afonine, P. V et al. Towards automated crystallographic structure refinement with phenix.refine. Acta Crystallogr. Sect. D 68, 352–367 (2012).

65. Adams, P. D. et al. PHENIX: A comprehensive Python-based system for macromolecular structure solution. Acta Crystallogr. Sect. D Biol. Crystallogr. 66, 213–221 (2010).

66. Schindelin, J. et al. Fiji: An open-source platform for biological-image analysis. Nat. Methods 9, 676–682 (2012).

