## supporting information for "Estrogen Receptor α/14-3-3 molecular glues as alternative treatment strategy for endocrine resistant breast cancer"

### Supplementary Figures and Tables

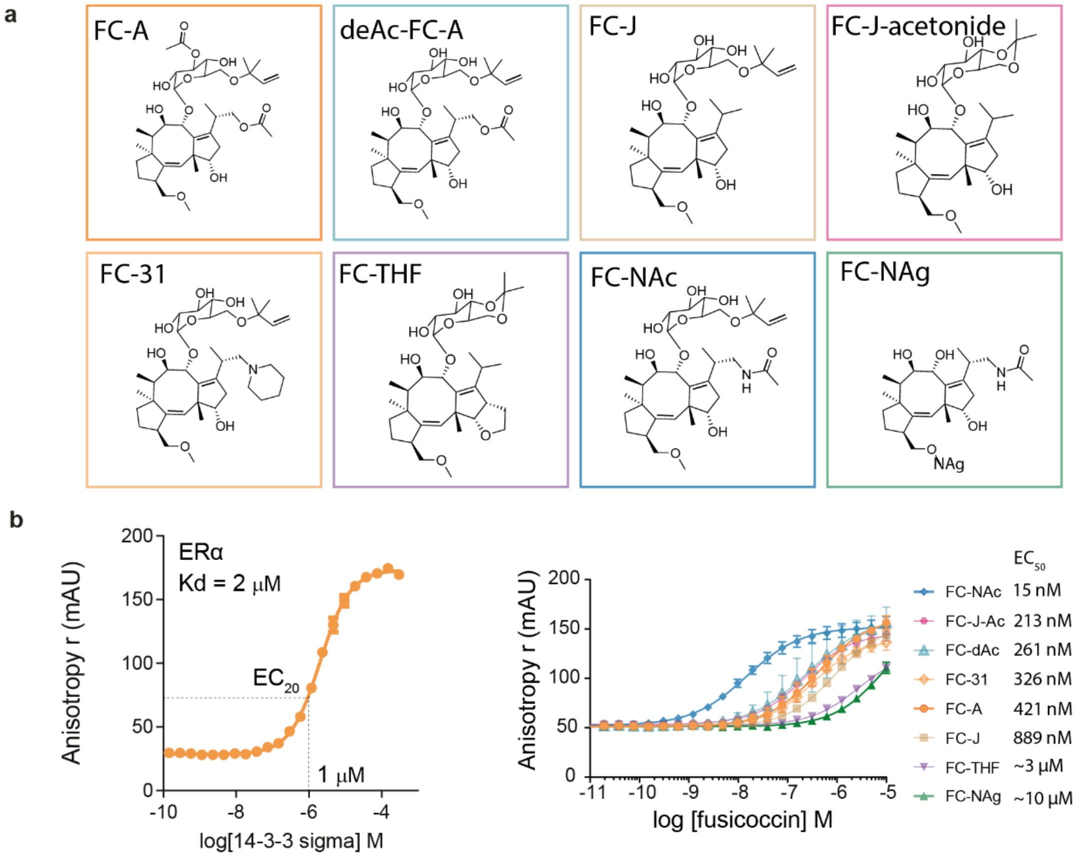


**Fig S1 | a**, Structure of FC-A and seven semi-synthetic derivatives. **b**, Left: Fluorescent Anisotropy assay of 14-3-3σ titrated to 10 nM FAM-labeled ERα-peptide. Right: Fluorescent Anisotropy assay of all fusicoccanes titrated to FAM-labeled ERα-peptide (10 nM) and 14-3-3σ (1 μM).

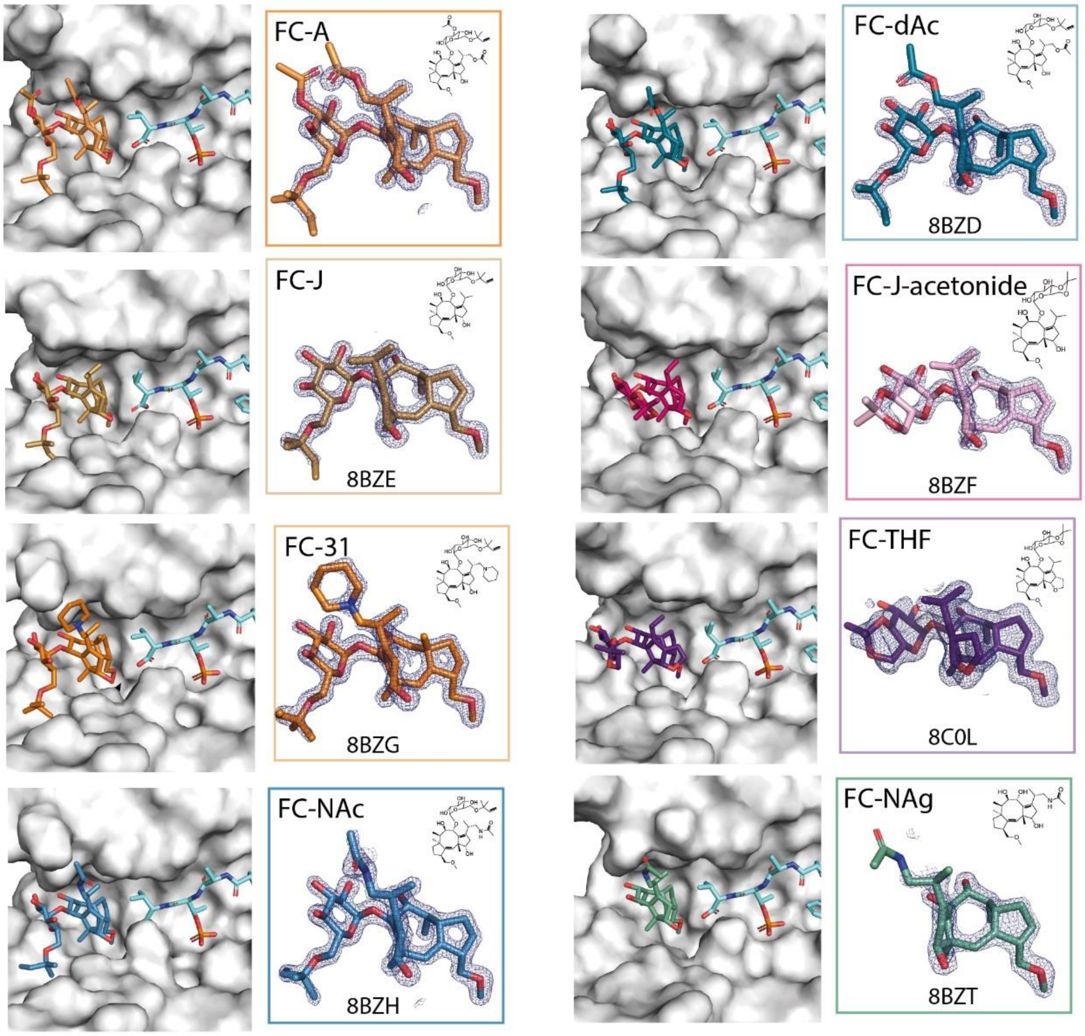


**Fig S2 |** Crystal structures and densities of FC-A and the seven semi-synthetic derivatives in complex with 14-3-3σ (white surface) and ERα (cyan sticks). 2Fo-Fc electron density maps (blue mesh) are contoured at 1σ. Crystallographic statistics are listed in table S1.

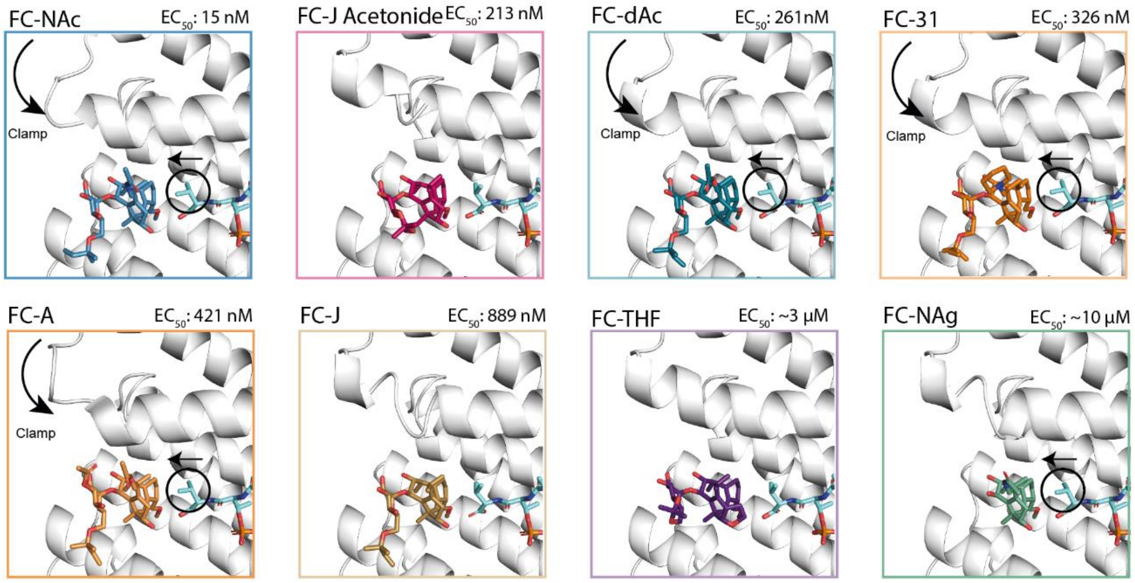


**Fig S3 |** Overview of helix-9 clamping effect of 14-3-3 (white cartoon) and conformational change of the C-terminal valine sidechain of ERα (cyan sticks) in the presence of the fusicoccanes.

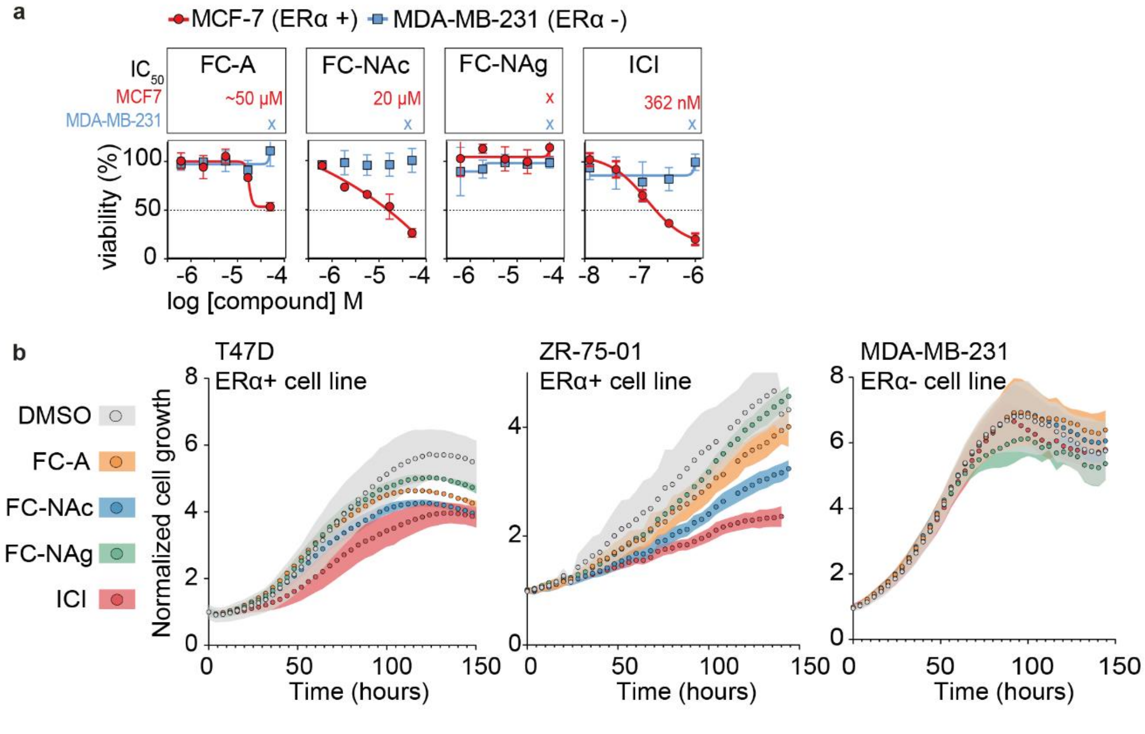


**Fig S4 | Effect on cell viability of fusicoccanes. a,** Cell titer blue viability assay of MCF-7 cells (red) and MDA-MB-231 cells (blue), treated with FC-A, FC-NAc, FC-NAg and ICI (mean ± SD, n = 3 biological independent). **b,** Cellular proliferation cells of T47D cells, ZR-75-01 cells and MDA-MB-231 cells, in the presence of DMSO (grey), 30 μM of FC-A (orange), 30 μM of FC-NAc (blue), 30 μM of FC-NAg (green), or 100 nM of ICI (red). (n = 3 biol. repl. with each 2 tech. repl.).

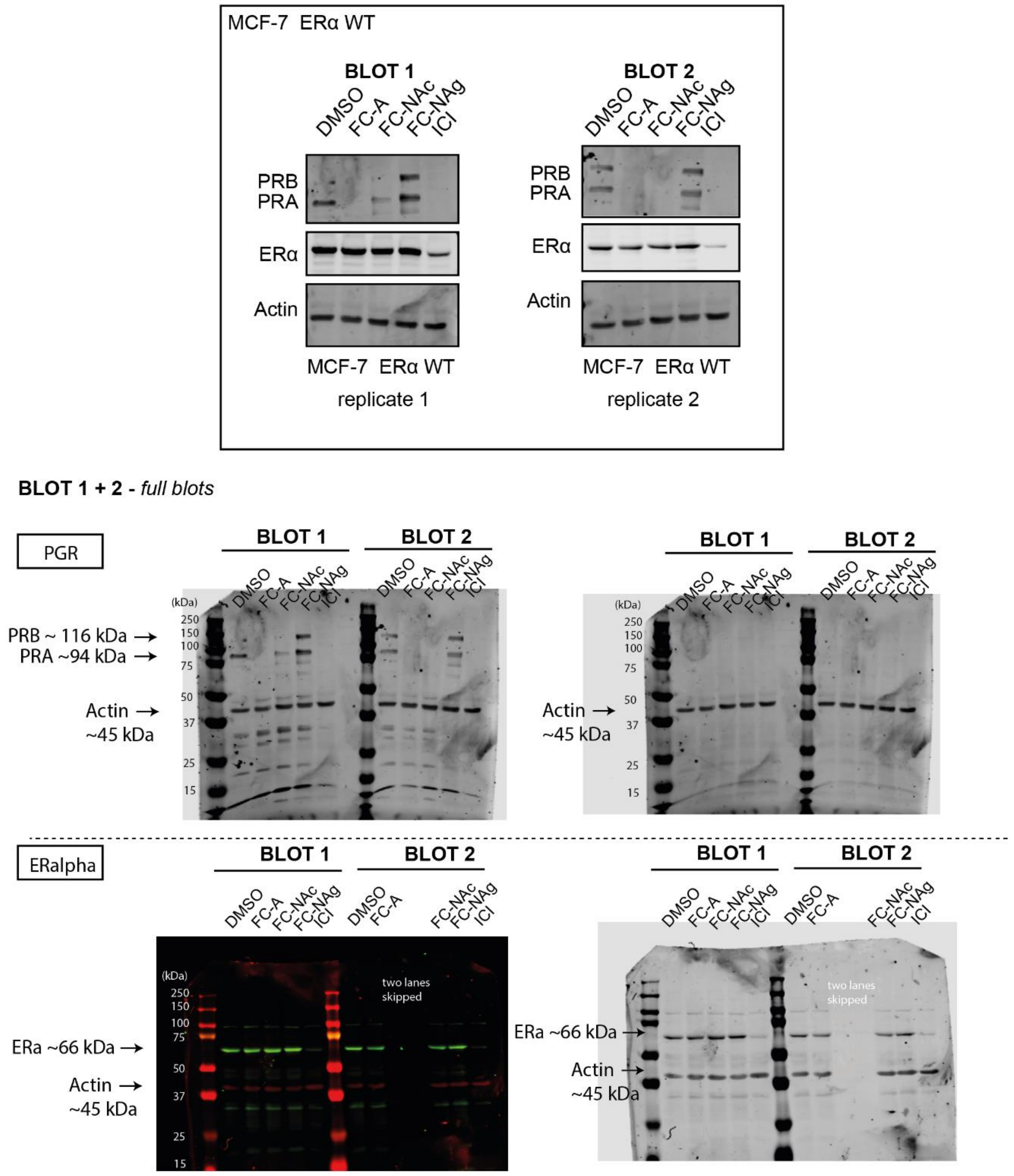


**Fig S5 |** Western blot replicates overview and full blots for down-stream protein (PR) of the ERα pathway for MCF-7 ERα WT cells upon treatment with FC-A, FC-NAc, FC-NAg (all 30 µM) or ICI (100 nM) in MCF-7 cells.

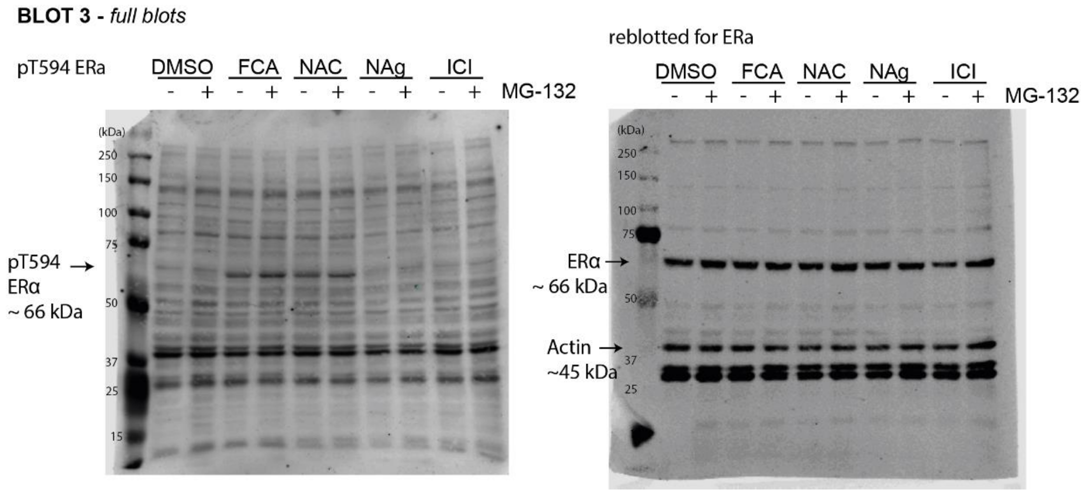


**Fig S6 |** unprocessed images for western BLOT 3 for (pT594) ERα.

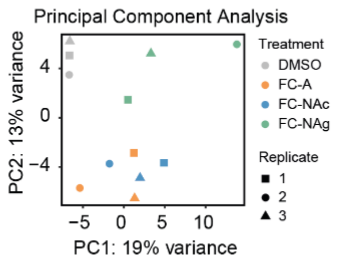


**Fig S7 |** Principal Component Analysis (PCA) of RNA-sequencing data.

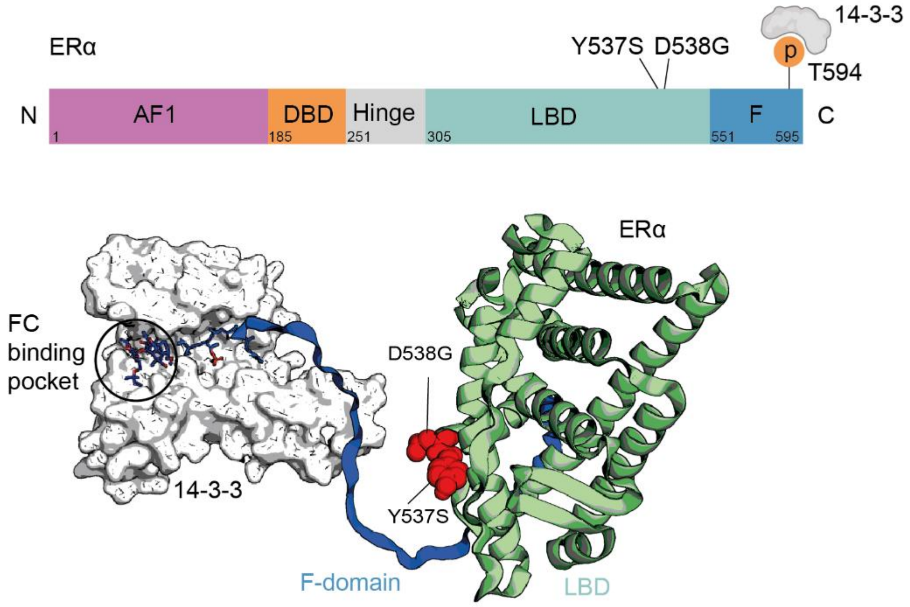


**Fig S8 |** Endocrine resistant patient mutations Y537S and D538G structurally don't interfere with ERα-14-3-3 stabilization.

**OVERVIEW BLOTS**

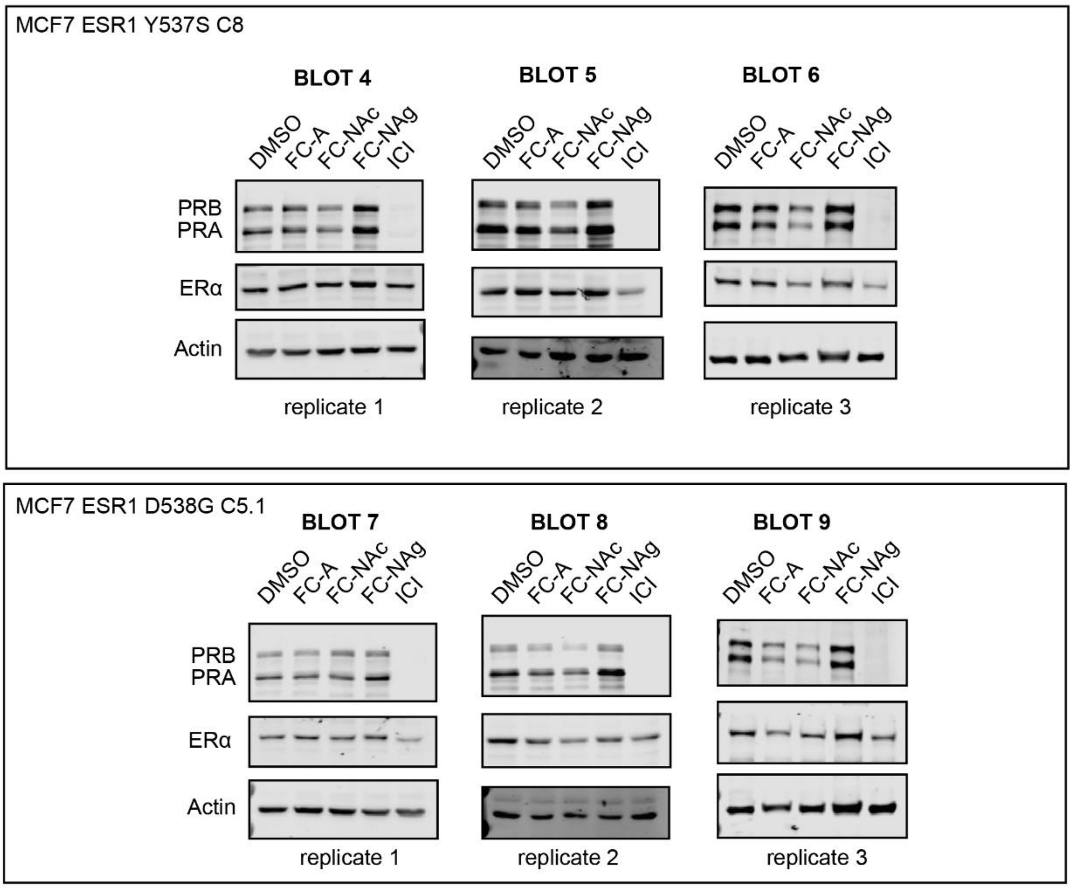


**Fig S9 |** Western blot overview replicates for down-stream protein (PR) of the ERα pathway for MCF-7 ERα Y537S (up) and D538G (down) cells.

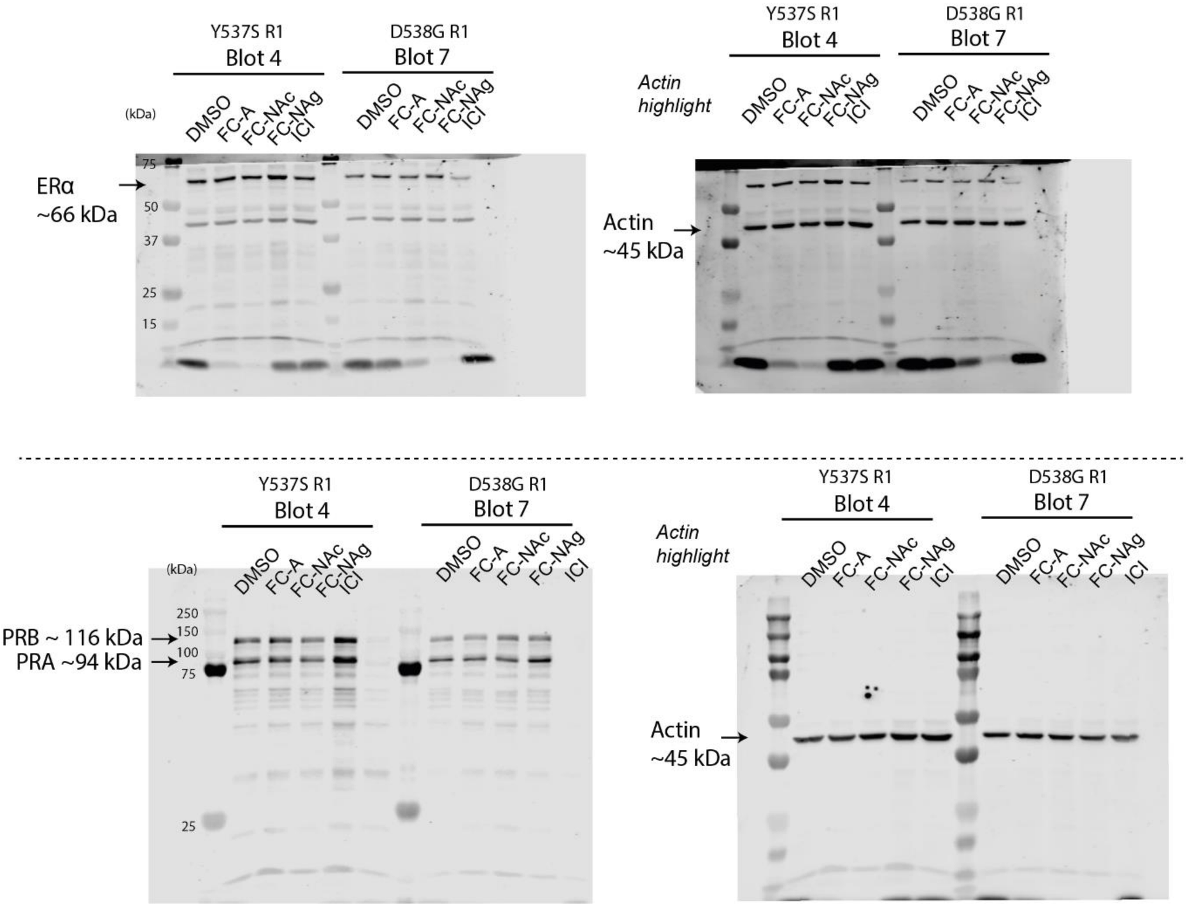


**Fig S10 |** unprocessed images for western blot 4 and 7 for replicate 1 of MCF7 ERα Y537S and D538G.

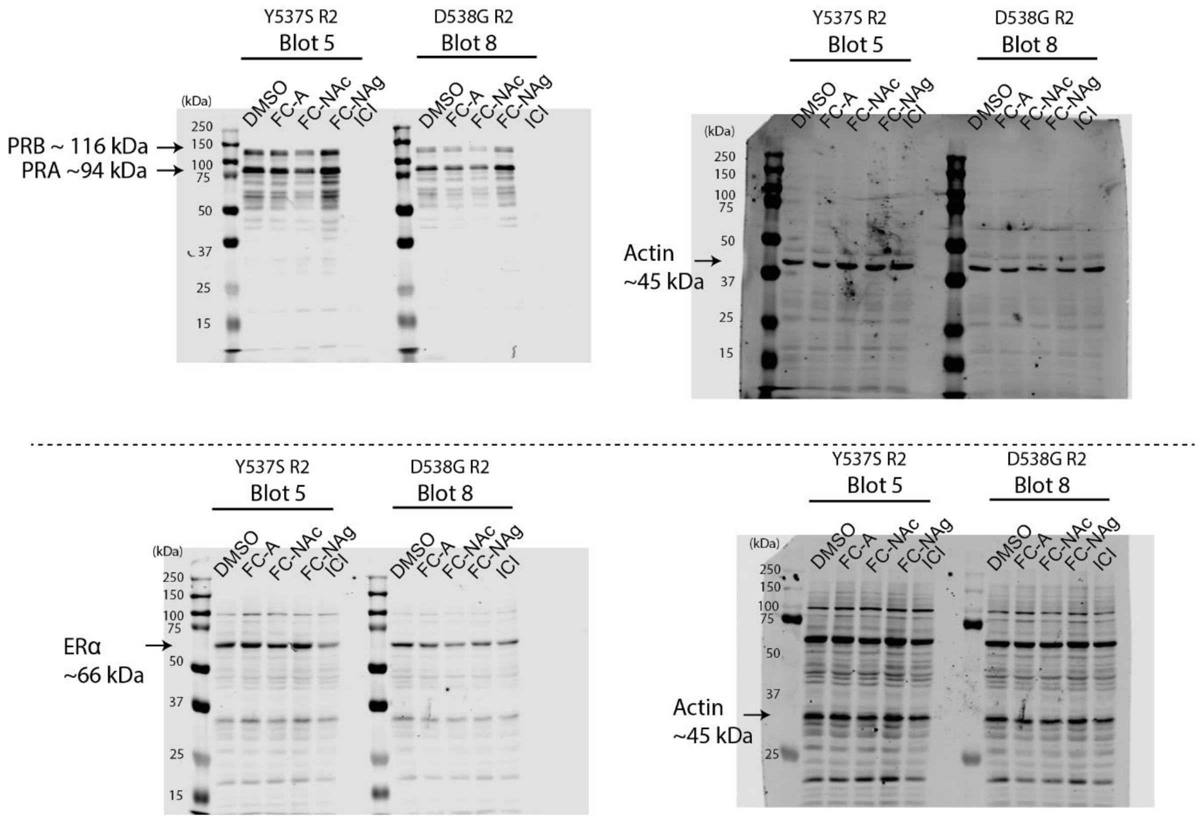


**Fig S11 |** unprocessed images for western Blot 5 and 8 for replicate 2 of MCF7 ERα Y537S and D538G.

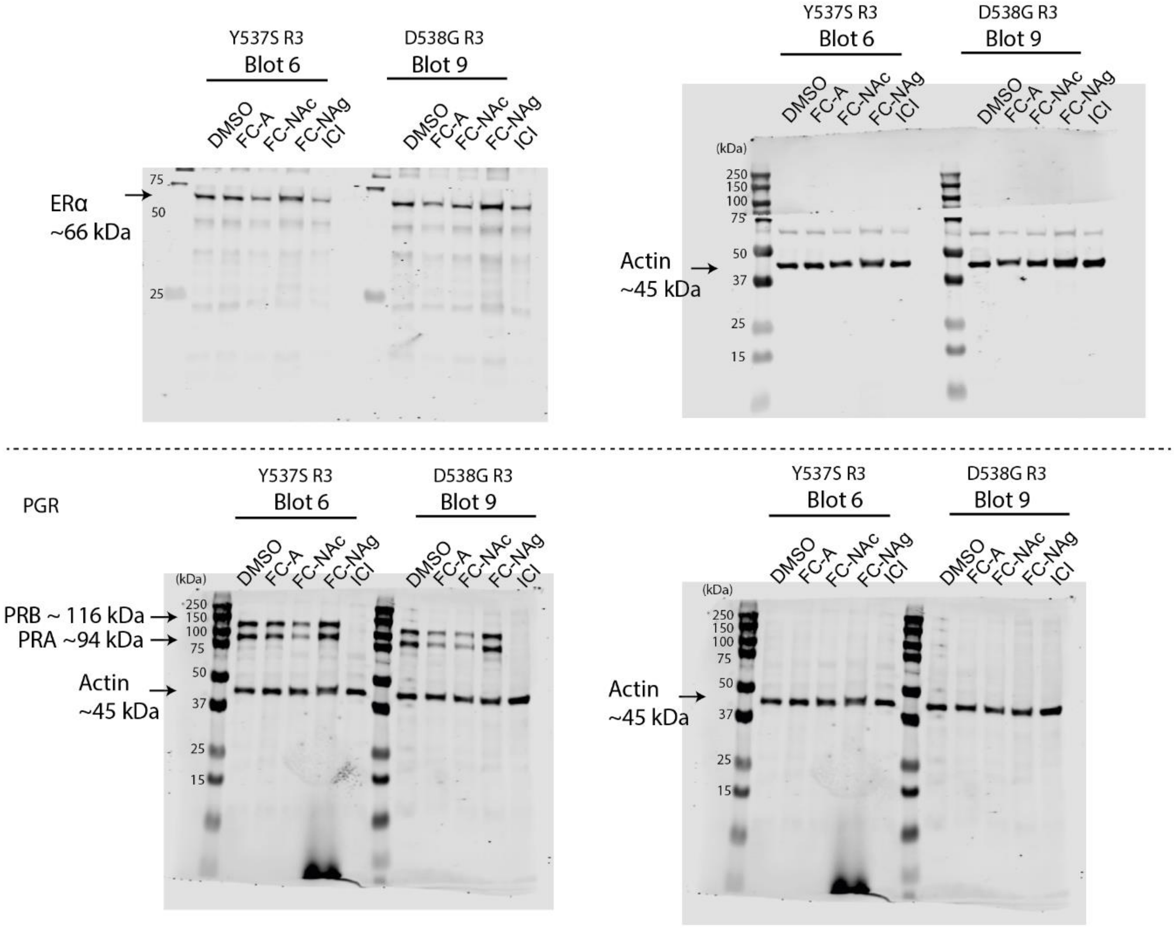


**Fig S12 |** unprocessed images for western blot 9 and 9 for replicate 3 of MCF7 ERα Y537S and D538G.

**BLOT 10**

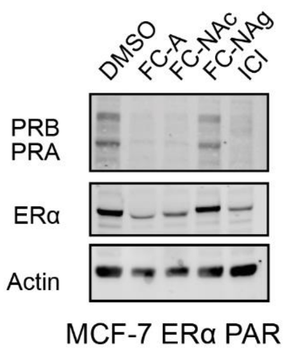


**BLOT 10**

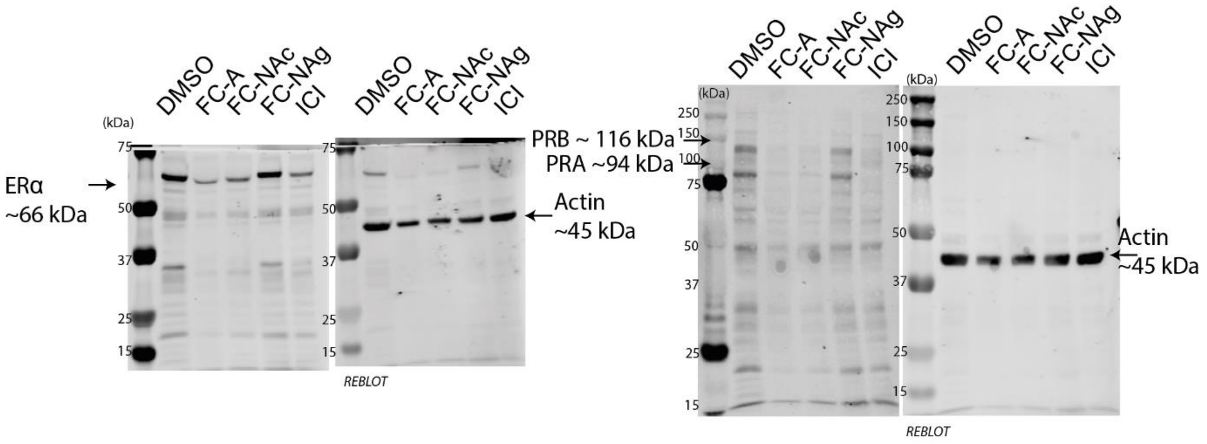


**Fig S13 |** Western blot of down-stream protein (PR) of the ERα pathway for MCF-7 parental cells treated with FC-A, FC-NAc, FC-NAg (all 30 µM) or ICI (100 nM). Down are the unprocessed images of blot 10.

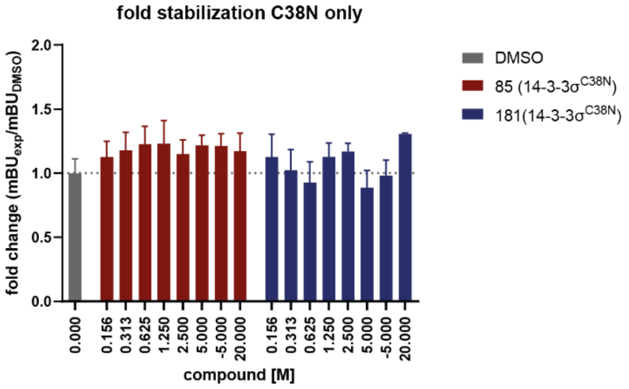


**Fig S14 |** NanoBRET signal observed for ERα/14-3-3σ C38N hybridization in HEK293T cells with increasing concentration of **85** (red) or **181** (blue), and DMSO (grey) (mean ± SD, n = 2).

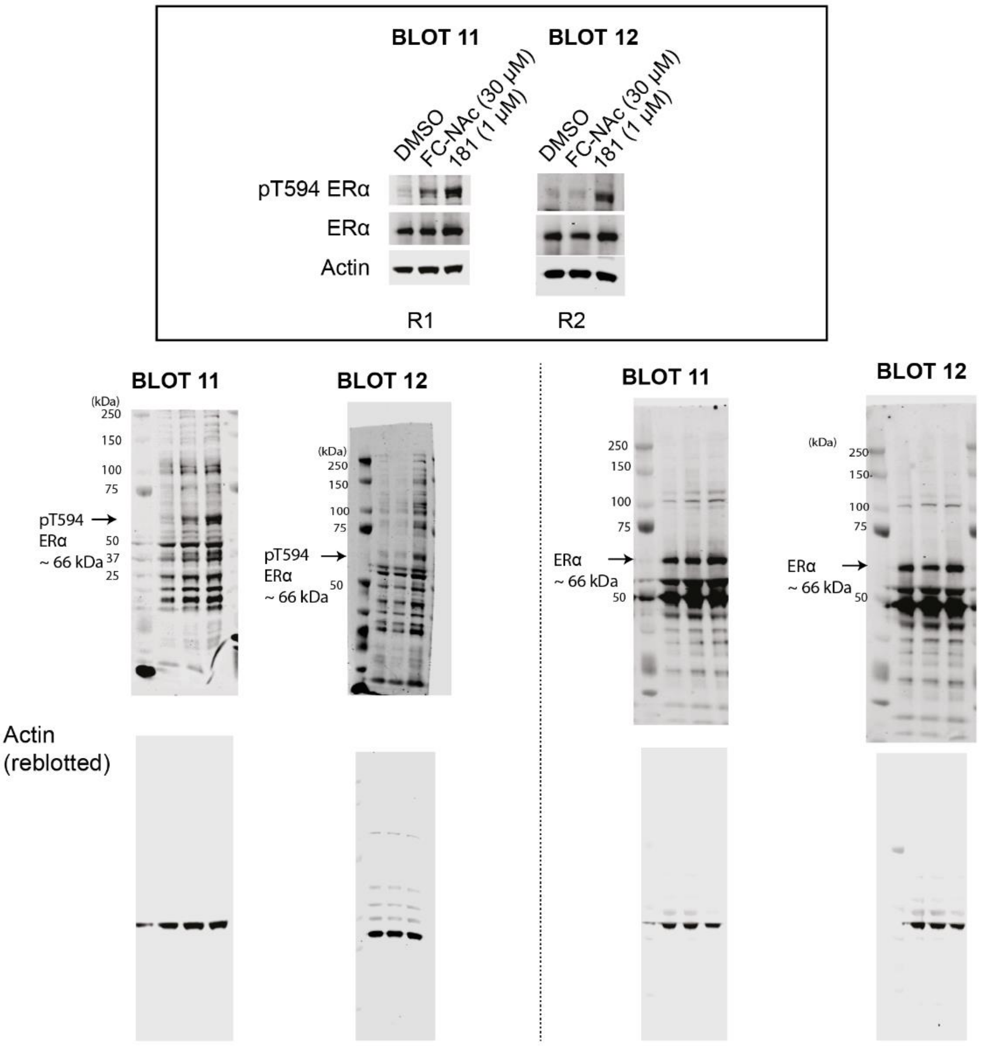


**Fig S15 |** Western blots and unprocessed images of MCF7 treated with DMSO, FC-NAc (30 µM) or **181** (1µM), stained for pT594-ERα and ERα protein levels.

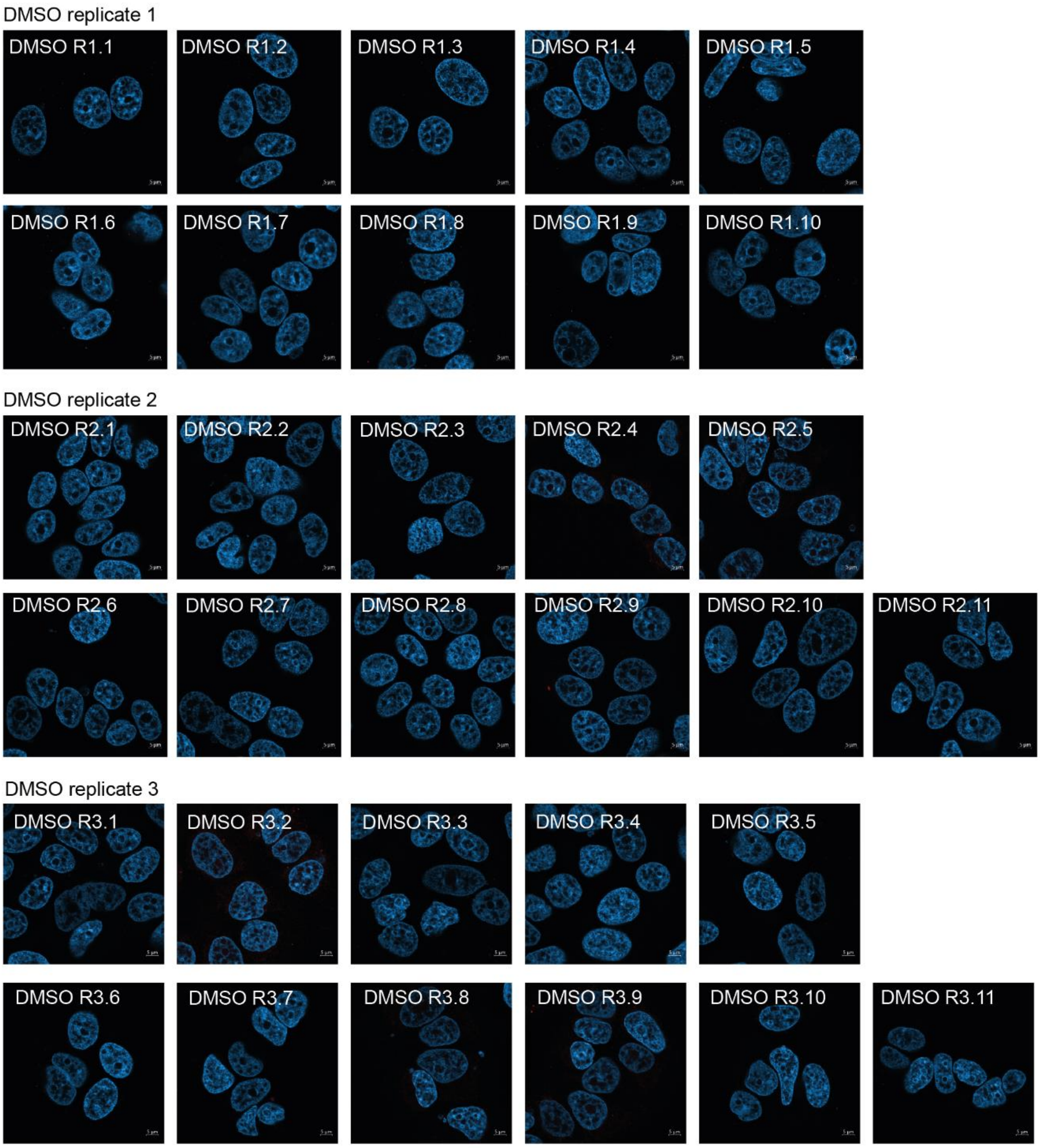


**Fig S16 |** Replicates of duolink proximity ligation assay for protein interactions between ERα and (pan) 14-3-3 in breast cancer cell line MCF-7, when treated with DMSO. Each red spot represents for a single interaction and DNA was stained with DAPI (blue).

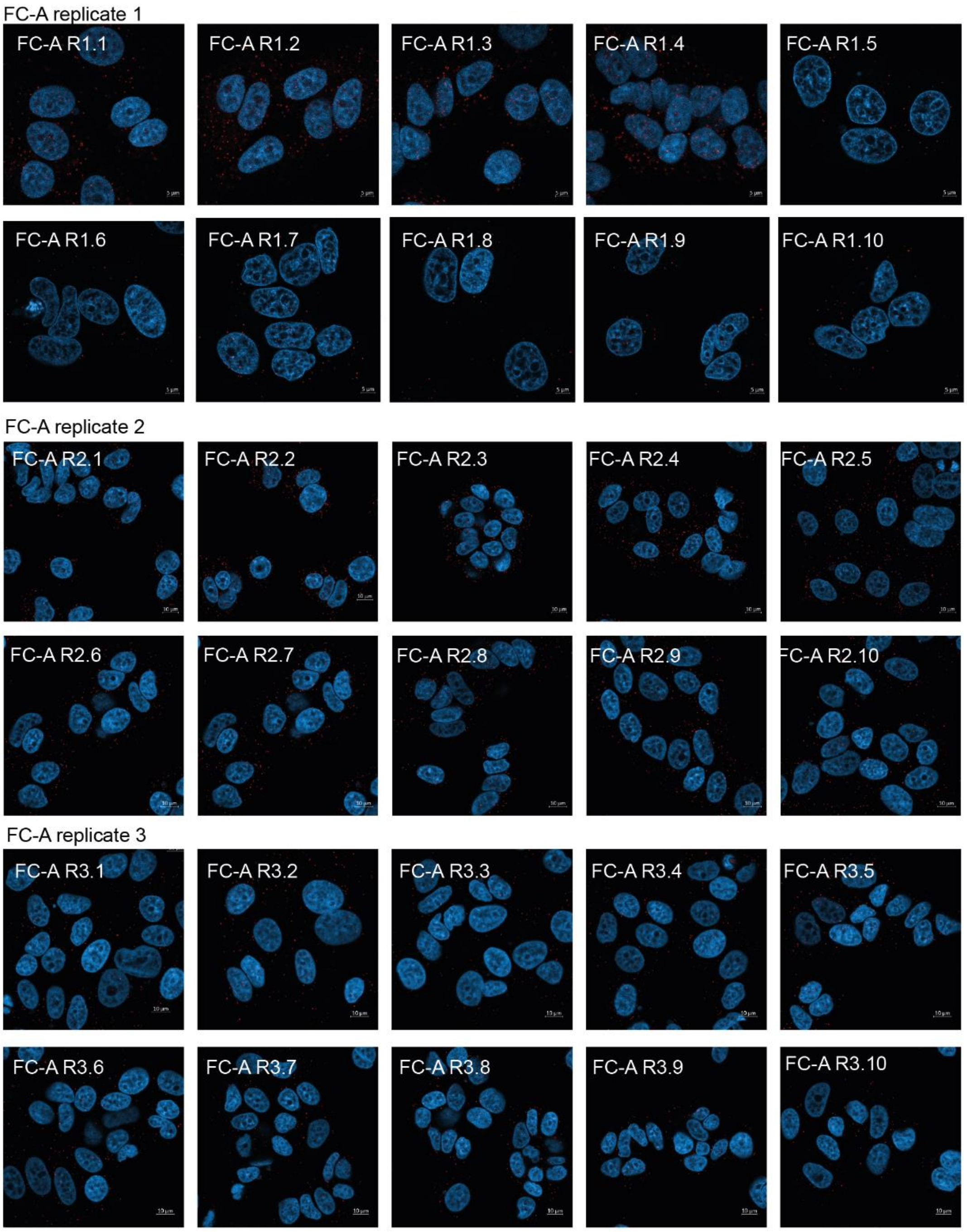


**Fig S17 |** Replicates of duolink proximity ligation assay for protein interactions between ERα and (pan) 14-3-3 in breast cancer cell line MCF-7, when treated with FCA (30 µM). Each red spot represents for a single interaction and DNA was stained with DAPI (blue).

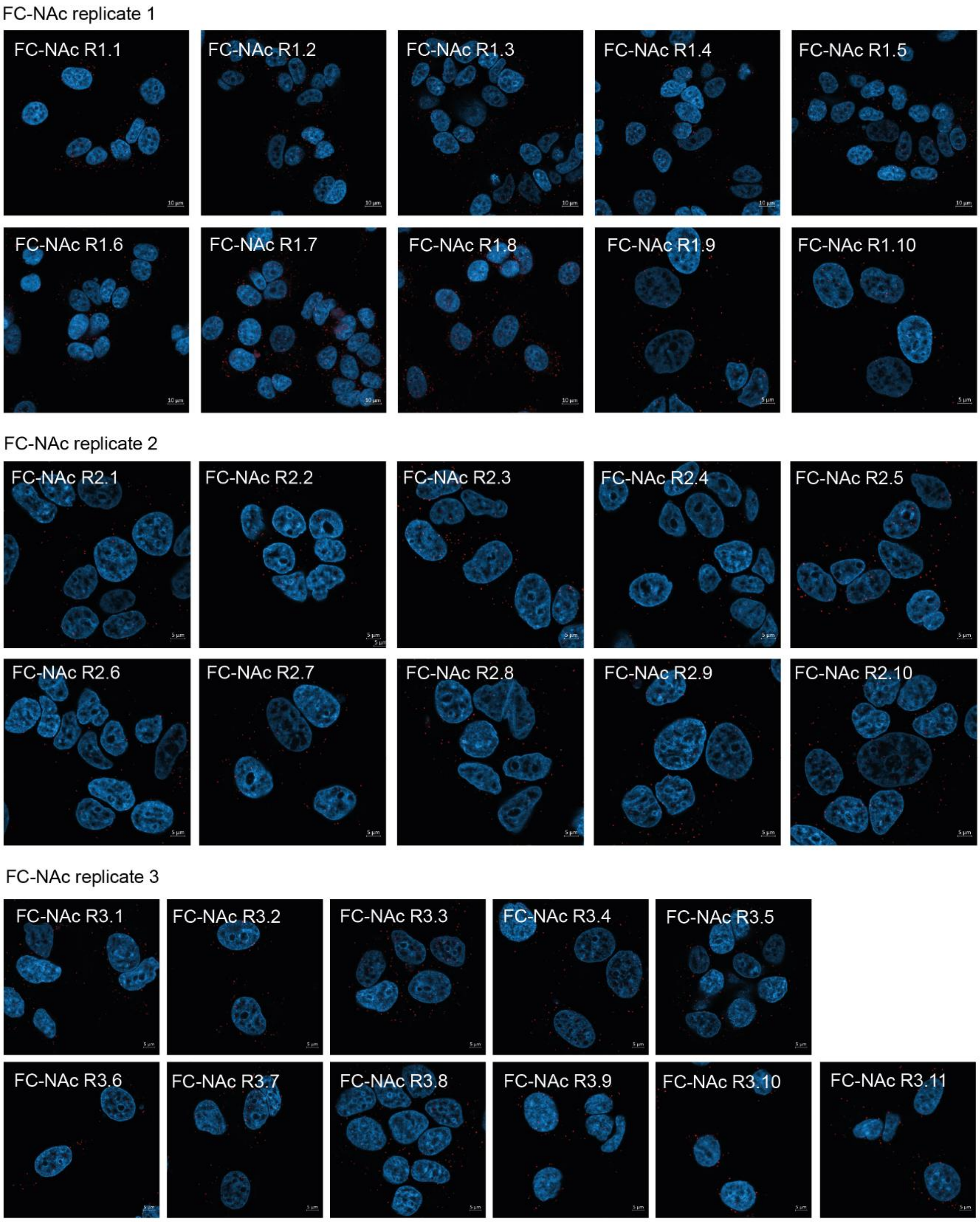


**Fig S18 |** Replicates of duolink proximity ligation assay for protein interactions between ERα and (pan) 14-3-3 in breast cancer cell line MCF-7, when treated with FC-NAc (30 µM). Each red spot represents for a single interaction and DNA was stained with DAPI (blue).

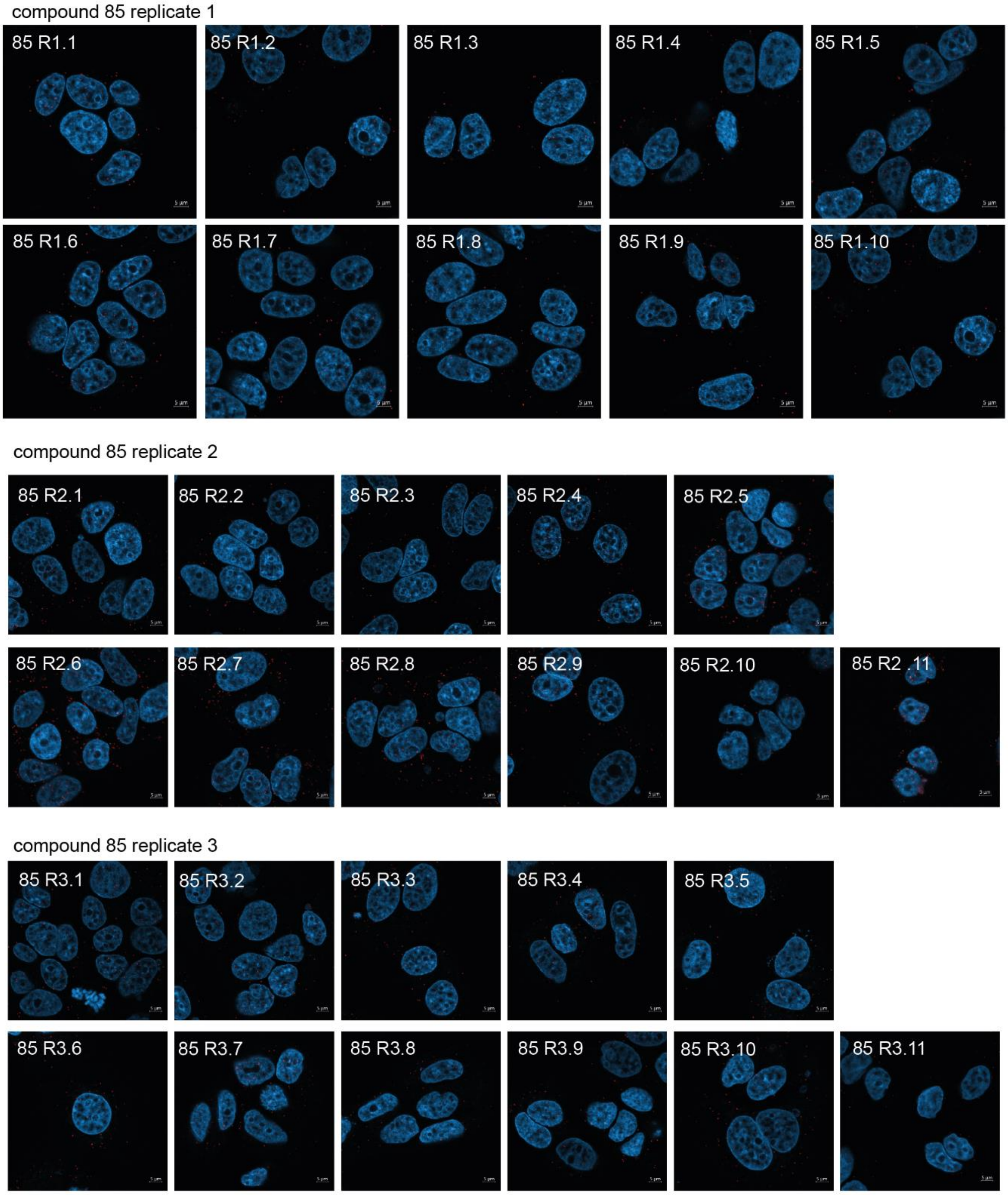


**Fig S19 |** Replicates of duolink proximity ligation assay for protein interactions between ERα and (pan) 14-3-3 in breast cancer cell line MCF-7, when treated with compound 85 (1 µM). Each red spot represents for a single interaction and DNA was stained with DAPI (blue).

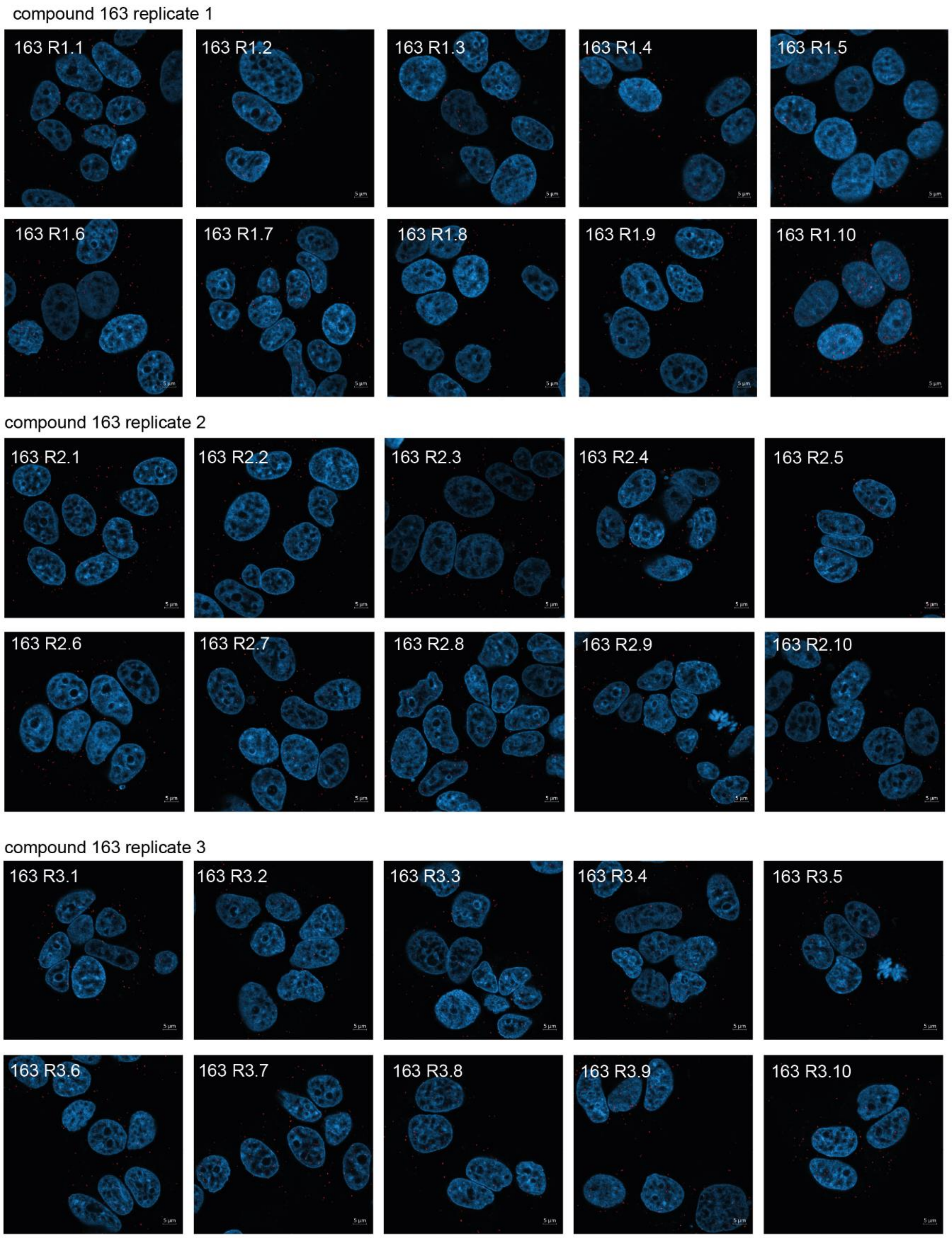


**Fig S20 |** Replicates of duolink proximity ligation assay for protein interactions between ERα and (pan) 14-3-3 in breast cancer cell line MCF-7, when treated with compound 163 (1 µM). Each red spot represents for a single interaction and DNA was stained with DAPI (blue).

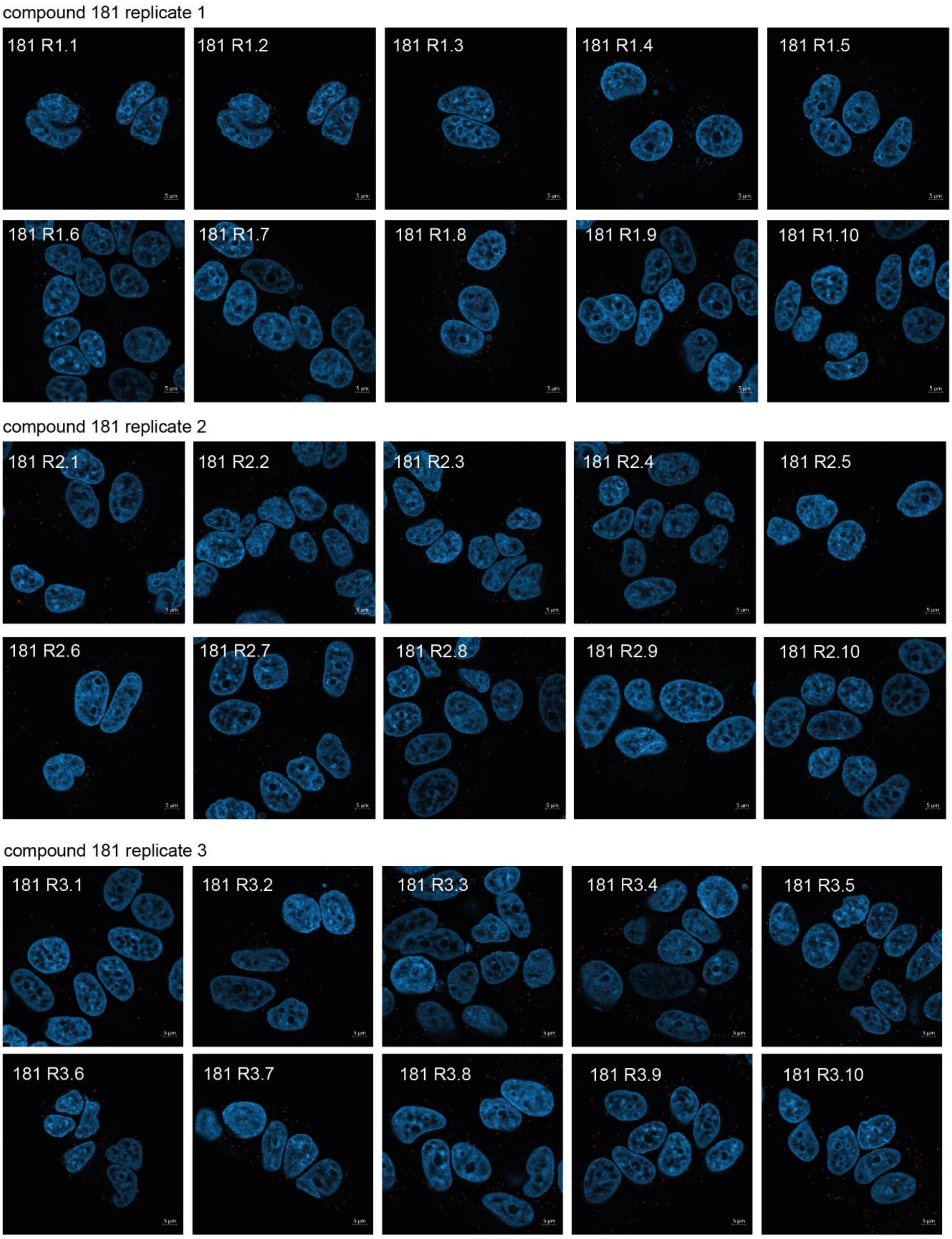


**Fig S21 |** Replicates of duolink proximity ligation assay for protein interactions between ERα and (pan) 14-3-3 in breast cancer cell line MCF-7, when treated with compound 181 (1 µM). Each red spot represents for a single interaction and DNA was stained with DAPI (blue).

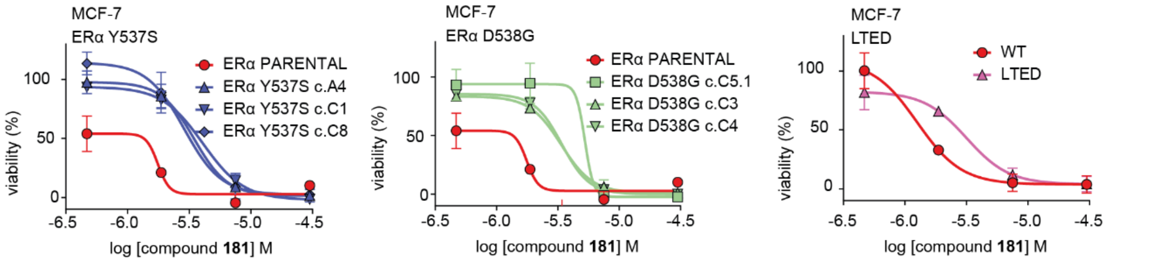


**Fig S22** | Cell titer blue (CTB) viability assay of **181** in MCF-7 ERα parental (PAR, red), ERα Y537S clones (blue, left), ERα D538G clones (green, middle), and long-term estrogen deprived (LTED) cells (pink, right). (mean ± SD, n = 3 biologically independent, with each 2 technical replicates).

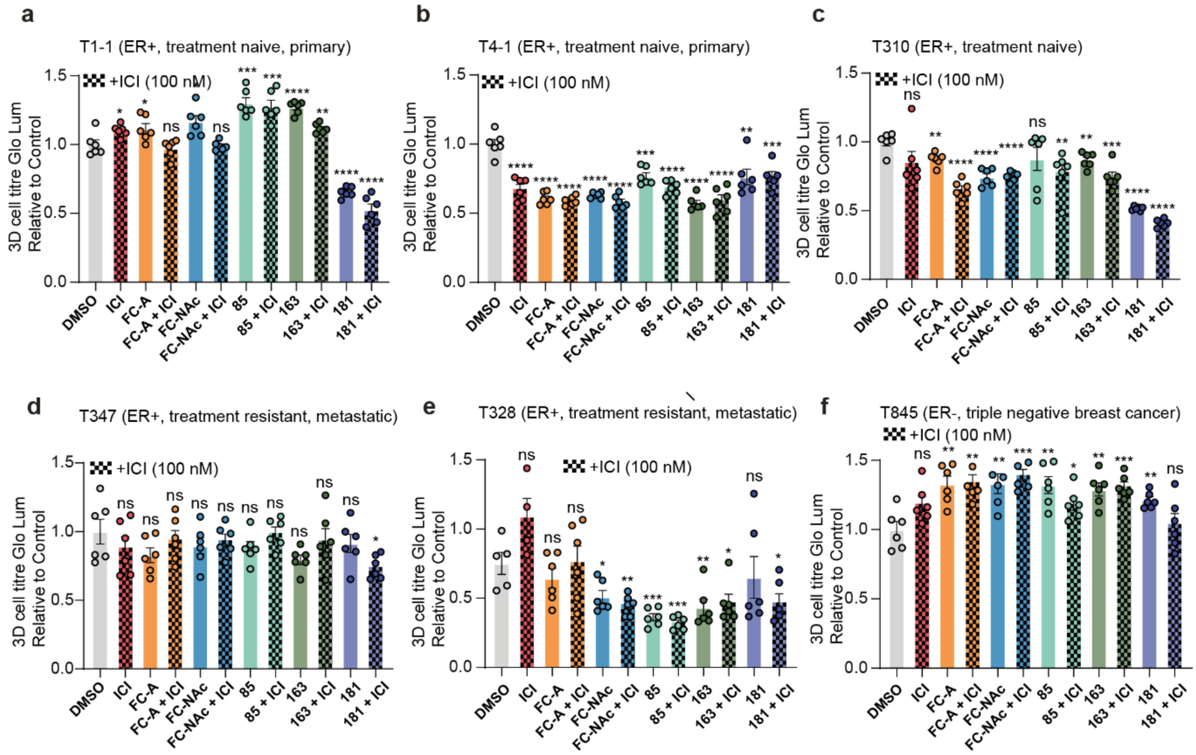


**Fig S23** | Bar charts of organoid culture proliferation subjected to vehicle control (DMSO), ICI (100 nM, pink), FC-A (30 µM, orange), FC-NAc (30 µM, blue),f compound **85** (1 µM, light green), compound **163** (1 µM, dark green) or compound **181** (1 µM, purple), all with and without ICI (100 nM, blocked bars). (mean ± SD, n = 4-6, biologically independent). (unpaired t-test: *p≤0.05, **p≤0.005 ***p≤0.001, ****p≤0.0001).

**Table S1: XRD data collection and refinement statistics for 14-3-3σ - ERα structures**

| 14-3-3σΔC / ERa | FC-dAc | FC-J | FC-J-acetonide | FC-31 | FC-THF | FC-NAc | FC-NAg |
|---|---|---|---|---|---|---|---|
| **PDB ID** | **8BZD** | **8BZE** | **8BZF** | **8BZG** | **8C0L** | **8BZH** | **8BZT** |
| **Data collection** | | | | | | | |
| **Collection source** | DESY PETRAIII P11 | DESY PETRAIII P11 | DESY PETRAIII P11 | DESY PETRAIII P11 | DESY PETRAIII P11 | DESY PETRAIII P11 | DESY PETRAIII P11 |
| **Collection date** | 07-05-2019 | 07-05-2019 | 07-05-2019 | 07-05-2019 | 09-06-2020 | 07-05-2019 | 07-05-2019 |
| **Wavelength (Å)** | 1.033200 | 1.033200 | 1.033200 | 1.033200 | 1.033200 | 1.033200 | 1.033200 |
| **Resolution (Å)** | 1.50 – 66.18 (1.50 – 1.53) | 1.43 – 66.15 (1.43 – 1.45) | 1.53 – 66.10 (1.53 – 1.56) | 1.49 – 66.15 (1.52 – 1.49) | 1.60 – 66.13 (1.60 – 1.63) | 1.46 – 66.12 (1.46 – 1.49) | 1.65 – 65.84 (1.65 – 1.68) |
| **Space group** | C 2 2 21 | C 2 2 21 | C 2 2 21 | C 2 2 21 | | C 2 2 21 | |
| **Unit cell** | 82.10 111.81 62.53 | 81.971 111.992 62.431 | 81.850 112.086 62.430 | 82.097 111.677 62.546 | 81.88 112.14 62.71 | 82.102 111.554 62.578 | 81.561 111.572 62.495 |
| **Total reflections**[a] | 590532 (17707) | 621844 (10703) | 565956 (21784) | 597109 (17456) | 497708 (22314) | 614743 (14242) | 446201 (19868) |
| **Unique reflections**[a] | 46188 (2098) | 52605 (2216) | 43668 (2142) | 47237 (2244) | 38541 (1854) | 49269 (2092) | 33913 (1669) |
| **Redundancy**[a] | 12.8 (8.4) | 11.8 (4.8) | 13.0 (10.2) | 12.6 (7.8) | 12.9 (12.0) | 12.5 (6.8) | 13.2 (11.9) |
| **Completeness (%)**[a] | 99.6 (92.7) | 98.6 (85.2) | 100.0 (100.0) | 99.9 (98.3) | 99.9 (97.9) | 98.2 (82.5) | 97.9 (100.0) |
| **Average I/σ(I)** [a] | 16.4 (2.7) | 22.6 (2.2) | 24.3 (1.7) | 24.0 (2.2) | 17.0 (1.3) | 23.6 (2.2) | 15.1 (2.2) |
| **Wilson B-factor ($Å^2$)** | 17.82 | 16.20 | 20.64 | 19.76 | 18.39 | 19.16 | 22.50 |
| **$CC_{1/2}$**[a,b,c] | 0.998 (0.854) | 0.999 (0.730) | 1.000 (0.692) | 0.999 (0.809) | 0.989 (0.862) | 0.999 (0.840) | 0.999 (0.820) |
| **$R_{merge}$**[a,c,e] | 0.084 (0.706) | 0.059 (0.621) | 0.062 (1.371) | 0.053 (0.842) | 0.091 (0.789) | 0.054 (0.739) | 0.093 (1.140) |
| **$R_{meas}$**[a,d,e] | 0.088 (0.753) | 0.062 (0.698) | 0.064 (1.444) | 0.055 (0.903) | 0.095 (0.824) | 0.056 (0.799) | 0.097 (1.191) |
| **Refinement** | | | | | | | |
| **Reflections in set: Refinement / R-free** | 46159 (4374) / 2000 (190) | 52570 (4640) / 1326 (122) | 43643 (4311) / 1998 (197) | 47210 (4614) / 1164 (114) | 38455 (3814) / 1970 (189) | 49245 (4202) / 2429 (221) | 33884 (3393) / 1686 (175) |
| **Non-H atoms:** | | | | 2225 / 283 | 2211 / 279 | 2223 / 283 | 2201 / 277 |
| **Overall / solvent** | 2256 / 316 | 2305 / 351 | 2225 / 291 | | | | |
| **$R_{work}$ / $R_{free}$ (%)** | 0.1807 (0.2344) / 0.2033 (0.2664) | 0.1738 (0.2425) / 0.1896 (0.2676) | 0.1826 (0.2452) / 0.2115 (0.2876) | 0.1799 (0.2376) / 0.1891 (0.2304) | 0.1800 (0.2676) / 0.2057 (0.3033) | 0.1744 (0.2789) / 0.1888 (0.2897) | 0.1831 (0.2854) / 0.2080 (0.3142) |
| **RMSD from ideal geometry:** | | | | | | | |
| **Bond length (Å) / angles (°)** | 0.010 / 1.04 | 0.009 / 1.05 | 0.010 / 1.15 | 0.010 / 1.04 | 0.009 / 0.97 | 0.009 / 1.05 | 0.014 / 1.17 |
| **Average protein B-factor ($Å^2$)** | 24.96 | 21.67 | 26.76 | 27.31 | 21.59 | 26.82 | 27.72 |
| **Ramachandran:** | | | | | | | |
| **Favored / outlier (%)** | 97.45 / 0.00 | 97.97 / 0.43 | 97.02 / 0.43 | 98.72 / 0.00 | 98.30 / 0.00 | 98.30 / 0.00 | 97.02 / 0.00 |
| **Clashscore** | 2.89 | 4.18 | 2.90 | 4.20 | 2.64 | 2.63 | 5.28 |

[a] Number in parentheses is for the highest resolution shell used in the refinement

[b] $CC_{1/2}$ = Pearson's intra-dataset correlation coefficient.

[c] $R_{merge}$ (= $R_{sym}$) = $\sum_h \sum_1 |I_{h1} - < I_h >| / \sum_h \sum_1 < I_h >$, where $I_{h1}$ is the intensity of the 1th observation of reflection h and $< I_h >$ is the average intensity of reflection h

[d] $R_{meas} = \sum_h \left| \sqrt{(n_h/(n_h - 1)} \sum_1 |I_{h1} - < I_h >| \right| / \sum_h \sum_1 < I_h >$ where $n_h$ is the number of observations of reflection $h$

[e] Correlation of experimental intensities with intensities calculated from refined model.
